# Rapid evolution of bacterial mutualism in the plant rhizosphere

**DOI:** 10.1101/2020.12.07.414607

**Authors:** Erqin Li, Ronnie de Jonge, Chen Liu, Henan Jiang, Ville-Petri Friman, Corné M.J. Pieterse, Peter A.H.M. Bakker, Alexandre Jousset

## Abstract

Even though beneficial plant-microbe interactions are commonly observed in nature, direct evidence for the evolution of bacterial mutualism in the rhizosphere remains elusive. Here we use experimental evolution to causally show that initially plant-antagonistic *Pseudomonas protegens* bacterium evolves into mutualists in the rhizosphere of *Arabidopsis thaliana* within six plant growth cycles (6 months). This evolutionary transition was accompanied with increased mutualist fitness via two mechanisms: *i)* improved competitiveness for root exudates and *ii)* enhanced capacity for activating the root-specific transcription factor gene *MYB72*, which triggers the production of plant-secreted scopoletin antimicrobial for which the mutualists evolved relatively higher tolerance to. Genetically, mutualism was predominantly associated with different mutations in the GacS/GacA two-component regulator system, which conferred high fitness benefits only in the presence of plants. Together, our results show that bacteria can rapidly evolve along the parasitism-mutualism continuum in the plant rhizosphere at an agriculturally relevant evolutionary timescale.

## Introduction

Mutualistic interactions between multicellular hosts and their associated microbiota are important for the fitness of both parties ^1–4^. However, while commonly observed in nature, direct evidence for the evolution of mutualism at both phenotypic and genotypic level is still scarce ^5–7^. The rhizosphere is a hotspot for mutualistic interactions between the plant and free-living microorganisms. For example, plants can preferentially interact with mutualistic microbes present in the indigenous species pool of the soil and disproportionally increase their relative abundances in the rhizosphere ^8–10^. While such plant-mediated ecological filtering can rapidly change the relative abundances of mutualistic versus antagonistic species in the rhizosphere, it is less clear if plants can drive evolution of mutualism within species by increasing the fitness of emerging *de novo* mutualist genotypes. For example, even the most well-known plant mutualistic microbes, nitrogen-fixing rhizobia ^5^ and phosphorus-providing mycorrhizae ^11^, can be detrimental to the plant, suggesting that the interaction between a given pair of plant and microorganism varies naturally ^12,13^. It is thus possible that plant-associated microbes might evolve along the parasitism-mutualism continuum in response to selection exerted by plants.

Beneficial symbioses between eukaryotic and prokaryotic organisms have evolved multiple times across the eukaryotic domain ^14^ and are considered as one of the major evolutionary transitions of life ^15^. It has been suggested that the evolution of mutualism often requires two basic components: currency and mechanism of exchange of the currency ^14^. In the context of plant-bacteria interactions, currency could be, for example, a root exudate, which can be taken up by bacteria. Similarly, bacteria might produce plant growth-promoting hormones such as auxin and gibberellins ^16^, that are beneficial for plant growth. When the currency exchange between both parties is symmetrical, the selection is expected to favour the evolution of mutualism. Increased mutualistic dependence is then thought to evolve via reciprocal coevolution or via adaptation by one of the partners via selection on traits that are directly involved in the mutualistic interaction ^15^. However, currency exchange could also be asymmetrical, due to competition for shared limiting nutrients, such as iron ^17^, which could explain why certain plant-microbe interactions are antagonistic. Moreover, due to the open nature of the rhizosphere, free diffusion of plant-derived resources could select for increased levels of cheating where mutant bacterial genotypes take advantage of ‘public goods’ without contributing to the production of plant growth-promoting compounds ^5,17^. As a result, mutualistic plant-microbe interactions might require additional enforcing from the plant ^5^ via sanctioning of cheating bacterial genotypes.

To assess whether plant-microbe mutualism can emerge as a consequence of plant-mediated effects, we used an *in vivo* experimental evolution design ^18^ where we allowed the rhizosphere bacterium *Pseudomonas protegens* CHA0 to evolve on the roots of *Arabidopsis thaliana* in the absence of other microbes. Furthermore, we used sterile sand free of organic carbon as the growth substrate making bacterial growth obligately dependent on plant root exudates. As a result, bacterial survival and evolution was solely dependent on the presence of the plant, and the performance of evolved bacterial selection lines were thus compared with the ancestral bacterial strain. To set up the selection experiment, we inoculated a clonal bacterial population on the roots of five independent *A. thaliana* Col-0 replicate plant selection lines and grew the plants and *P. protegens* in otherwise gnotobiotic conditions for a total of six plant growth cycles, which lasted four weeks each. At the end of every growth cycle, the evolved bacterial populations were isolated and transferred to the rhizosphere of new sterile plants (Fig. S1). In these experimental conditions, the initial plant-bacterium interaction was antagonistic: *A. thaliana* aboveground biomass was clearly reduced in the presence of *P. protegens* CHA0 after one growth cycle (F_1, 8_ = 45.4, *P*< 0.001, Fig. 1A), and likely cause for this is the production of diverse bioactive metabolites by CHA0 ^19^ that can constrain plant growth ^20^. To quantify changes in plant-bacterium interaction, sixteen evolved bacterial colonies were randomly selected from each plant replicate selection line at the end of the second, fourth and sixth growth cycles, in addition to sixteen randomly selected ancestral colonies (in total 256 isolates). Each isolated colony was characterized phenotypically by measuring multiple key life-history traits, including growth on different carbon sources and media, tolerance to diverse abiotic and biotic stresses, production of several bioactive compounds and their ability to inhibit other microorganisms (Table S1). A subset of bacterial phenotypes was also full genome sequenced and characterised for their effects on plant growth in terms of root architecture and above and belowground biomasses at the end of the selection experiment.

**Figure 1.**
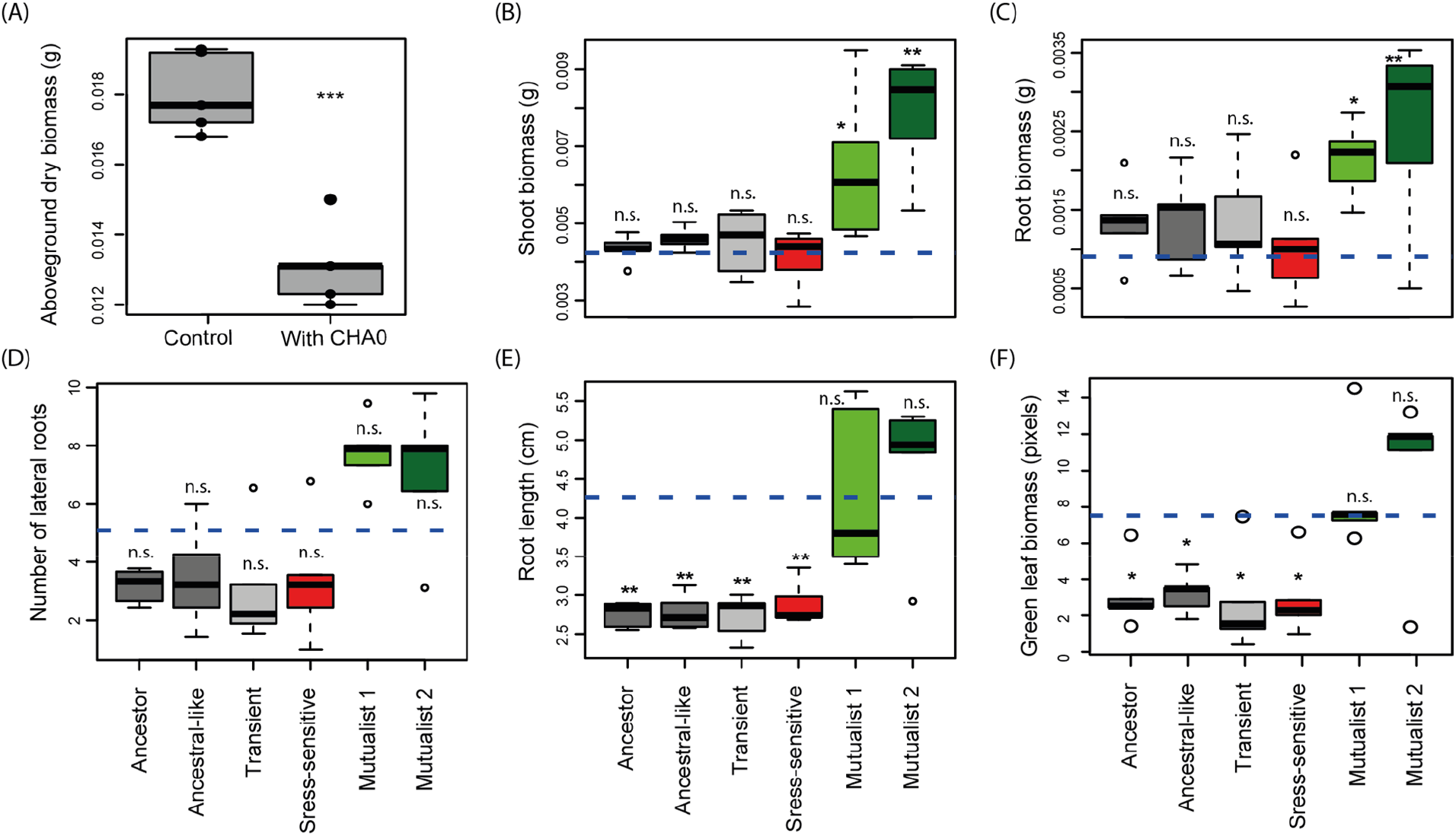
Evolution of bacterial mutualism in the rhizosphere of *Arabidopsis thaliana*. Panel (A) shows the initially antagonistic effect of *Pseudomonas protegens* CHA0 on *A. thaliana* after one plant growth cycle in the sterile sand study system. Panels (B-F) compare the effects of ancestral and evolved *Pseudomonas protegens* CHA0 phenotypes on plant performance-related traits in a separate plant growth assays performed on agar plates at the end of the selection experiment. Different panels show the shoot biomass in grams (B), root biomass in grams (C), number of lateral roots (D), root length in cm (E) and the amount of plant ‘greenness’ in terms of green-to-white pixel ratio (F) 14 days after bacterial inoculation (Supplementary dataset 2; blue dashed horizontal lines show the non-inoculated control plants). Bacterial phenotype groups are displayed as different colours and were classified and named based on K-means clustering (Figure S1) using 14 phenotypic traits linked to growth, stress tolerance, production of bioactive compounds, and antimicrobial activity (Table S1). Boxplots show the mean effect of five representative bacterial isolates from each evolved phenotype in addition to ancestor isolates (See Table S2). Statistical testing in all panels was carried out using ANOVA, and asterisks above plots indicate significant differences between control plants and bacteria-treated plants (*α=0.05, **α=0.01, ***α=0.001; n.s. = non-significant).

## Results

### Selection in the plant rhizosphere leads to bacterial phenotypic diversification and evolutionary transition towards mutualism

To study the evolution of *P. protegens* CHA0 in the *A. thaliana* rhizosphere, we isolated a total of 240 evolved bacterial isolates from every second time point along with sixteen ancestral isolates (Supplementary dataset 1) and used K-means clustering analysis to separate them into five distinct phenotypic groups based on measured all life-history traits (Fig. S2, Table S2). The phenotypic groups were then given names that reflected key differences in their appearance, life-history traits, and their mean effects on plant growth (Fig. 1; Fig. S3–S4). Evolved clones that clustered together with the ancestral strain were named as ‘Ancestral-like’ phenotype. Another phenotype similar to the ancestral strain, which only appeared momentarily before dropping below detection level was named as ‘Transient’ phenotype (Figs. 1–2). A third phenotype that resembled the ancestral strain, but which had clearly reduced abiotic stress tolerance (F_5, 248_ = 40.8, *P*< 0.001, Fig. S4) and increased ability to form a biofilm (F_5, 249_ = 196.8, *P*< 0.001, Fig. S4), was named as ‘Stress-sensitive’ phenotype. Agar plate assays were used to determine the effect of the evolved phenotypes on *A. thaliana* growth. While the ‘Ancestral-like’, ‘Transient’ and ‘Stress-sensitive’ phenotypes showed neutral effects on plant biomass relative to plant-only control lines (Shoot biomass, F_6, 26_ = 8.01, *P*< 0.001; Root biomass, F_6, 26_ = 2.84, *P*= 0.029, Fig. 1), they had a negative effect on the plant root length (Root length, F_6, 26_ = 10.01, *P*< 0.001, Fig. 1E) and resulted in clear bleaching of plants indicative of reduced chlorophyll activity similar to the ancestor (the amount of green pixels, F_6, 26_ = 5.90, *P*< 0.001, Fig. 1F). These assays also revealed two novel phenotypes that showed positive effects on plant shoot and root biomasses (Fig. 1B, C) with comparable levels of plant ‘greenness’ to no-bacteria control treatment (Fig. 1F). These evolved phenotypes were therefore named as ‘Mutualist 1’ and ‘Mutualist 2’as indicated by their plant growth-promoting activity.

**Figure 2.**
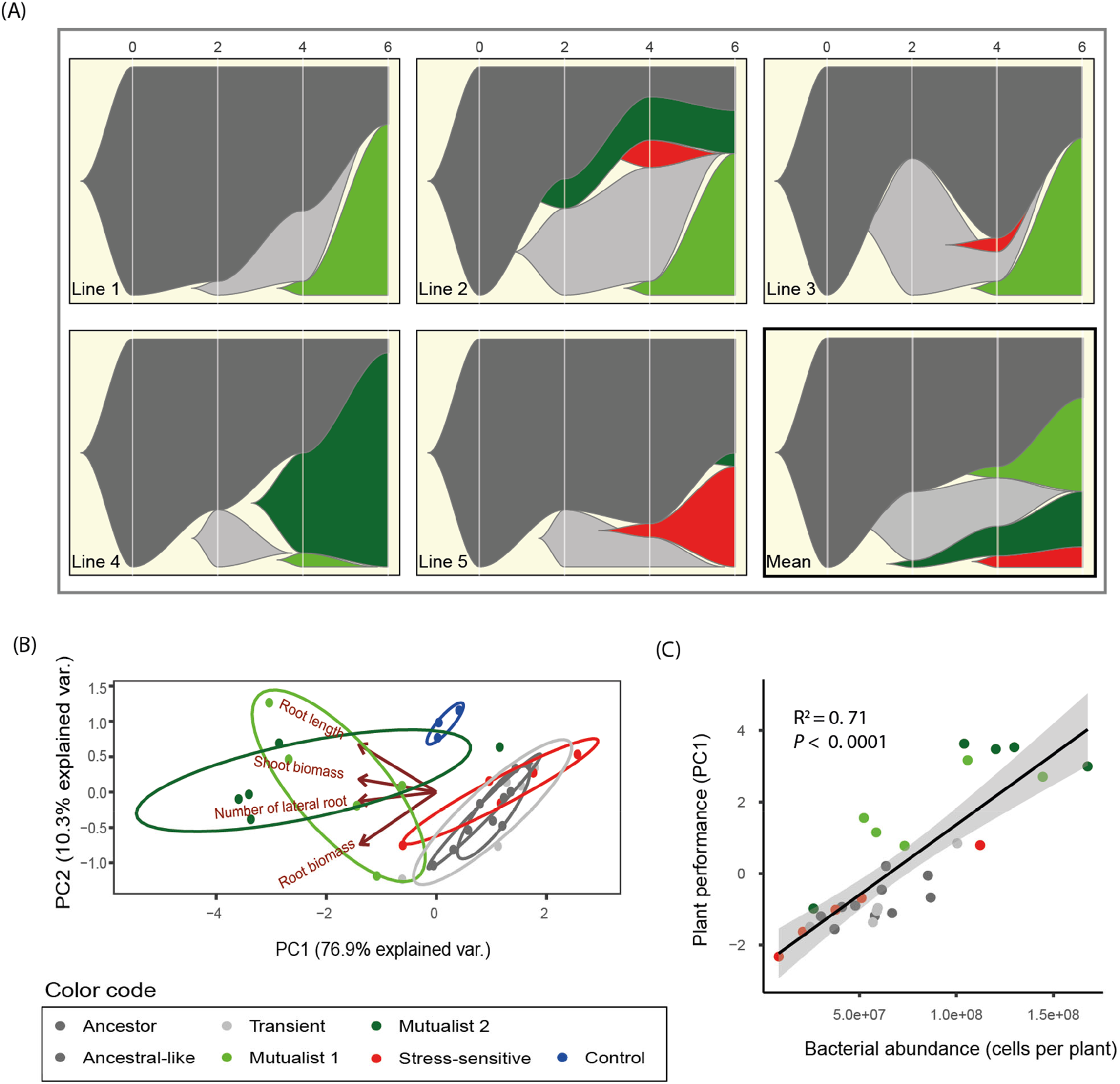
Temporal changes in bacterial phenotypes during the selection experiment and positive correlation between evolved bacteria and plant growth. Panels in A show the dynamics of five bacterial phenotype groups across five plant replicate lines and the overall mean pattern during six growth cycles (6 months). The x-axis shows the plant growth cycle (0: ancestral bacterium), and the y-axis shows the relative abundance of each bacterial phenotype. Panel (B) shows a principal component analysis (PCA) for five representative bacterial isolates from each evolved phenotype group in addition to ancestor isolates (See Table S2) based on their plant growth-related traits. The negative PC1 values of each isolate were extracted and combined to a ‘Plant performance’ index, which included bacterial effects on shoot biomass, root biomass and root architecture explaining 76.9% of the total variation in plant growth. Panel (C) shows a positive correlation between ‘Plant performance’ and bacterial abundance in the plant roots at the end of the fitness assays.

The relative abundance of different phenotype groups changed over time (Fig. 2A). The ‘Ancestral-like’ phenotypes persisted throughout the experiment even though they were substituted by evolved phenotypes in all plant selection lines (Fig. 2A). The evolutionary success of ‘Transient’ and ‘Stress-sensitive’ phenotypes was short-lived: ‘Transient’ phenotypes disappeared below the detection limit in all plant selection lines by the end of the sixth growth cycle (Fig. 2A), while the ‘Stress-sensitive’ phenotypes emerged only in three selection lines and survived until the end of the experiment only in one of the selection lines (Fig. 2A). In contrast, the frequency of mutualistic phenotypes increased in four out of five plant selection lines throughout the experiment, while one selection line became dominated by ‘Ancestral-like’ and ‘Stress-sensitive’ phenotypes (Fig. 2A).

An aggregated ‘plant performance’ index summarising the effects of each bacterial isolate on both aboveground and belowground plant growth traits (Fig. 2B, PC1 of multivariate analysis), was used to explore if reduced antagonism towards the plant was associated with improved bacterial growth indicative of the evolution of reciprocally beneficial mutualistic interaction. We found a significant positive correlation between plant performance index and bacterial phenotype abundance per plant (F_1, 28_ = 8.01, *P*< 0.001, Fig. 2C). Specifically, both mutualistic phenotypes reached higher abundances in the plant rhizosphere compared to other phenotypes (bacterial cells per plant). This indicates that reduced bacterial antagonism towards the plant was coupled with improved growth in the rhizosphere, which could also explain why mutualists became the dominant phenotypes in four out of five plant selection lines during the selection experiment (reaching up to 94% relative abundance, Fig. 2A). In support for this, a similar positive correlation was observed between the degree of plant performance of each phenotype measured in separate plant growth assays and their relative abundance in diversified rhizosphere populations at the end of the selection experiment (F_1, 23_ = 4.37, *P*= 0.048, Fig. S5). Together, these results demonstrate that the evolution of plant-growth promotion was accompanied with increased bacterial fitness, indicative of a mutualistic interaction where each species had a net benefit. As this evolutionary transition was observed parallel in four out of five selection lines, it was likely driven by deterministic processes such as selection exerted by the plant instead of random genetic drift due to bottlenecking between plant growth cycles.

### Evolution of mutualism is linked to improved resource catabolism and tolerance to plant-secreted antimicrobials

For stable mutualism to evolve, plants would need to provide the evolved mutualists a ‘currency’ that could not be accessed by the other phenotypes or employ some form of ‘sanctioning’ to constrain the growth of non-mutualist phenotypes. To study this, we first compared differences in the evolved phenotypes’ ability to use a range of carbon sources that are typically found in *A. thaliana* root exudates ^21^, and which could have selectively preferred the growth of mutualist phenotypes. Second, we compared the evolved phenotypes’ tolerance to scopoletin, which is an antimicrobial secreted by plant roots known to modulate root microbiome composition for example by favouring more tolerant bacterial taxa ^22,23^. We found that the ‘Mutualist 1’ phenotype showed an improved ability to grow on various different carbon sources compared to the other phenotypes (PC1 of multivariate analysis, F_5, 248_ = 50.72, *P*< 0.001, Fig. 3A). This suggests that ‘Mutualist 1’ potentially evolved an ability to compete better for plant-derived root exudates during the selection experiment, which could have increased their abundance relative to other phenotypes. Moreover, ‘Transient’, ‘Stress-sensitive’ and ‘Mutualist 2’ phenotypes showed reduced growth on carbon sources relative to ‘ancestral-like’ phenotypes indicative of competitive disadvantage (Fig. 3A). To explore the potential significance of scopoletin, we used a GUS reporter assay to determine how different bacterial phenotypes affected the expression of the plant root-specific gene *MYB72* ^24^, which encodes a transcription factor that regulates the production of scopoletin. We found that plants inoculated with ‘Mutualist 1’ and ‘Mutualist 2’ phenotypes retained high GUS activity, which was comparable to the ancestor (F_5, 24_ =5.6, *P* < 0.01, Fig. 3B, Fig. S6). However, the other evolved phenotypes induced a reduced GUS activity in plant relative to the mutualists (Fig. 3B, Fig. S6). While the ‘Mutualist 1’ and ‘Mutualist 2’ phenotype groups included strains that showed very high tolerance to scopoletin, their mean tolerance did not significantly differ from other phenotype groups (F_5, 24_ = 1.76, *P* = 0.16; Fig. 3C). However, a significant positive correlation was observed between the induction of *MYB72* in *A. thaliana* roots and phenotypes’ tolerance to scopoletin (F_1, 28_ =8.29, *P* < 0.01, Fig. 3D). This suggest that both the activation of scopoletin production, and the scopoletin tolerance, were potentially under co-selection as mutualistic phenotypes showed high scopoletin tolerance only when they were able to trigger increased scopoletin production by the plant (upregulation of *MYB72*). Such mechanisms would have ensured positive selection for mutualists relative to other evolved phenotypes.

**Figure 3.**
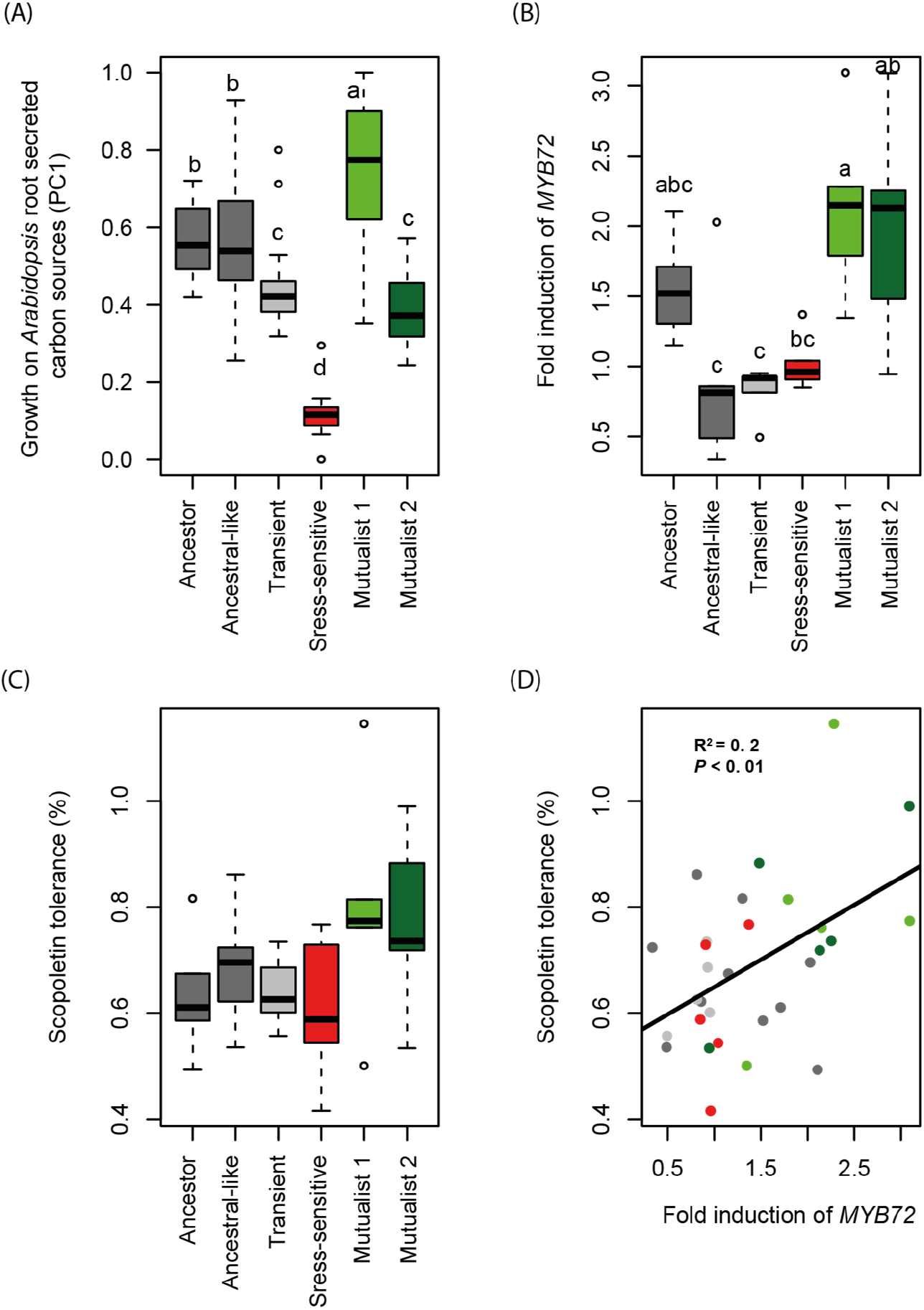
Selection mechanisms favouring the increase in the relative abundance of mutualists in the rhizosphere of *Arabidopsis thaliana*. Panel (A) shows the growth of ancestor and evolved *Pseudomonas protegens* CHA0 phenotypes on carbons typically secreted by *A. thaliana* (14 most dominant carbons analysed as a combined index based on normalised first principal component (PC1, which explained 83.9 % of total variation); in total, 16 ancestral, 119 ‘Ancestral-like’, 11 ‘Stress-sensitive’, 37 ‘Mutualist 1’, 31 ‘Mutualist 2’ and 41 ‘Transient’ isolates were characterized, Supplementary dataset 1). Panel (B) shows the effect of ancestor and evolved *P. protegens* CHA0 phenotypes on the expression of *MYB72* (transcription factor responsible for scopoletin production) in the roots of a GUS *A*. *thaliana* reporter line (based on the quantification of GUS staining of the roots, Fig. S6). Panel (C) shows the relative growth of ancestor and evolved *P. protegens* CHA0 phenotypes in the presence of the plant-secreted scopoletin antimicrobial at 2 mM concentration after 96 hrs of incubation relative to no-scopoletin control. Panel (D) shows a positive relationship between *MYB72* expression (fold induction; x-axis) and scopoletin tolerance (y-axis) for all tested isolates. Panels (B-D) include five representative bacterial isolates from each phenotype in addition to the ancestor (each replicate line represented; See Table S2). In all panels, colours represent different phenotype groups and statistical testing in panels (A-C) was carried out using ANOVA (different letters indicate significant differences based on a Tukey HSD test (α=0.05)).

### Mutualistic phenotypes had mutations in genes encoding the GacS/GacA two-component regulatory system

To gain insights into the genetic mechanisms underlying the evolution of mutualism, we performed whole-genome re-sequencing of a subset of evolved isolates followed by reference-based identification of point mutations (SNPs) and insertions or deletions (INDELs). These analyses revealed that different evolved bacterial phenotypes were associated with relatively few mutations in global regulator genes (Fig. 4) underpinning their central role in bacterial adaptation ^25,26^. While only few non-parallel mutations were observed in case of ‘Ancestral-like’ and ‘Transient’ phenotypes (Table S2), all but two mutualistic isolates (8/10) harboured mutations in genes encoding the GacS/GacA two-component system, which regulates secondary metabolism alongside many other aspects of bacterial physiology ^27^. Despite high level of parallelism, a variety of different mutations was observed in this locus. Three *gacS*/*gacA* mutations were unique to ‘Mutualist 1’ isolates, and specifically associated with an uncharacterized N-terminal histidine kinase domain in GacS (G27D) and the response regulatory domain of GacA (G97S; D49Y, Fig. 4). Four unique ‘Mutualist 2’ mutations were found upstream (at −40 of the transcription start site) or inside the *gacA* coding region (E38*, D54Y and Y183S, Fig. 4). The conserved phosphate-accepting aspartate 54 (D54) residue is important for phospho-relay initiated by the sensor kinase GacS, and mutations of this residue are associated with complete loss-of-function ^28–30^. Aspartate 49 (D49) is another conserved residue in the vicinity of D54, and the *gacA* (49Y) allele has previously been reported to be associated with a partial reduction in GacA activity ^30^. The other mutations in *gacA* are novel and conceivably have a significant impact on GacA activity as they result in a severely truncated protein (E38*) or are located within the third recognition helix of the LuxR-like tetra-helical helix-turn-helix (HTH) domain, which is known to make most of the DNA contacts (Y183S) ^31^. In line with the predicted effects of the mutations, ‘Mutualist 1’ isolates retained part of the GacS/GacA – mediated traits, while ‘Mutualist 2’ isolates showed a severe to complete disruption of secondary metabolite production (Fig. S3). These differences between individual mutations are thus likely to explain the variation in life-history traits within and between the ‘Mutualist 1’ and ‘Mutualist 2’ phenotype groups.

**Figure 4.**
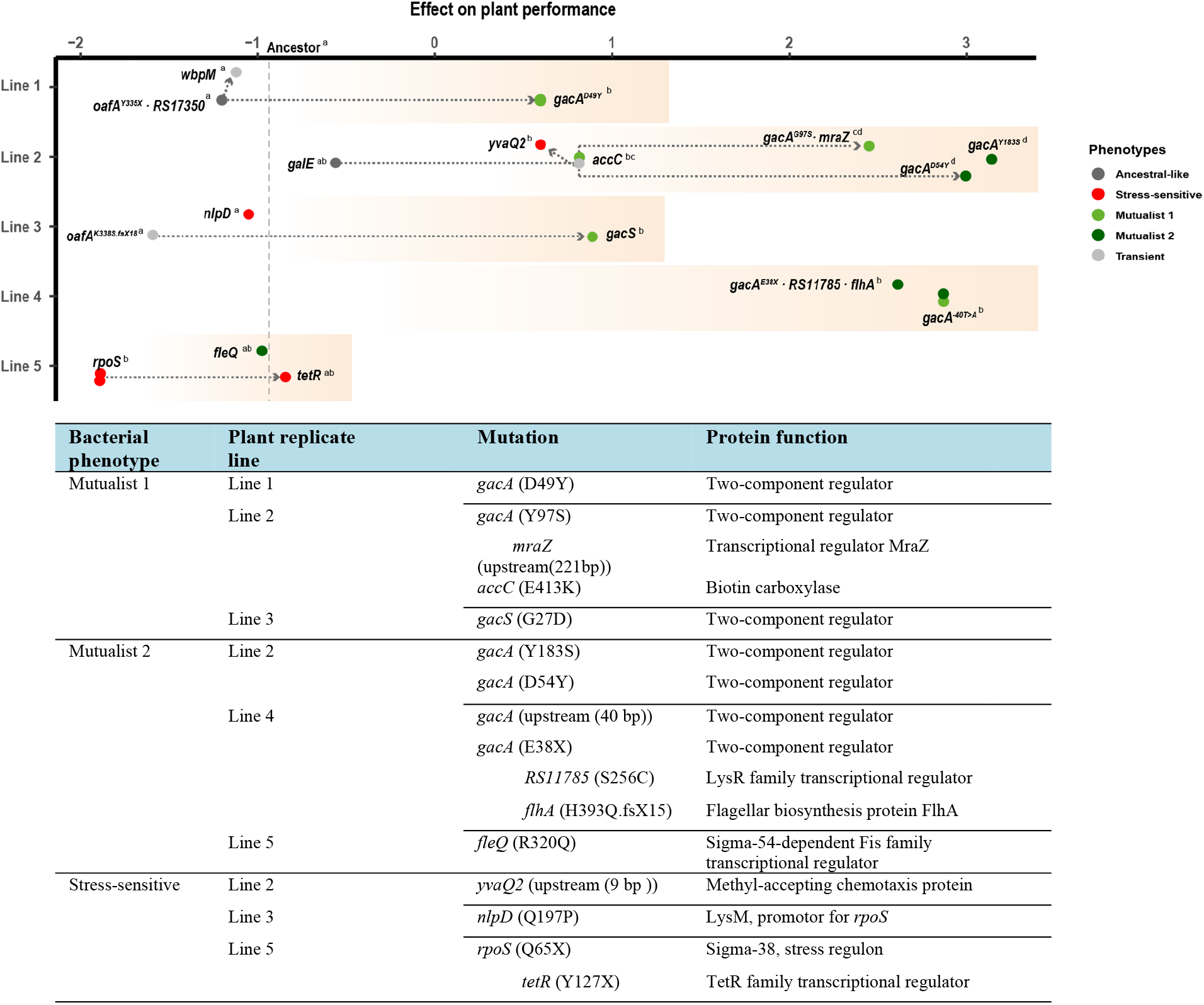
Genetic basis of bacterial evolution in the rhizosphere of *Arabidopsis thaliana*. Clear parallel evolution was observed between four out five plant replicate selection lines based on re-sequencing of 25 evolved and five ancestor isolates used in the phenotypic assays. Filled dots represent isolates with non-synonymous mutations (present in 18/25 evolved isolates), and the x-axis shows a combined index of ‘Plant performance’ relative to non-inoculated control plants (values on the x-axis indicate positive and negative effects on the plant and the y-axis shows the five independent plant replicate selection lines). The effect of the ancestral bacterial genotype on plant performance is shown as a vertical dashed line. The different letters on the top right of each genotype indicate significant differences based on a Tukey HSD test (α=0.05; each line analysed separately). The accumulation of mutations within replicate lines are shown with connected dashed arrows. The table lists unique mutations (and the strains’ ID number) linked with evolved bacterial phenotypes, and additional mutations that appeared later during the experiment within the same genetic background are shown after the indent; notably, these additional mutations did not affect the bacterial phenotypes (See Table S3 for a more detailed description of the mutations).

Several other genes were also mutated in the mutualistic isolates, including *accC* (E413K) that encodes the biotin decarboxylase subunit of the acetyl coenzyme A carboxylase complex involved in fatty acid biosynthesis in bacteria, and *fleQ* (R320Q) which is linked with motility and biofilm formation (Fig. 4). These mutations were plant replicate line-specific, and their effect on bacterial physiology or bacteria-plant interactions are unknown. Interestingly, mutualists evolved in all except one selection line, which became dominated by ‘Stress-sensitive’ bacteria (Fig. 1, Fig. 4), that also transiently appeared in two other plant selection lines. Genetically, this phenotype was mainly associated with mutations in the *rpoS* coding region and its promoter (Q65*, 3/5 of selection lines). The *rpoS* gene encodes sigma factor *sigma-38*, which mediates general stress resistance ^32,33^, downregulates the biosynthesis of antagonistic secondary metabolites ^34^ and is involved in biofilm formation ^35^ with several bacteria. In line with this, we found that ‘Stress-sensitive’ phenotypes were able to form high amounts of biofilm *in vitro* (Fig. S4), which may have supported more efficient root colonization ^36^ and explain their dominance in one of the plant selection lines. Moreover, efficient root colonisation could have initiated a strong priority effect ^37,38^, potentially constraining the subsequent emergence of mutualistic bacterial phenotypes. One of the sequenced ‘Stress-sensitive’ clones had also a mutation in a TetR-family transcriptional regulator (*tetR*), indicative of generalised stress tolerance evolution. Together, these results show that plant selection can lead to high level of parallel evolution both at the phenotypic and molecular level.

### The fitness benefits of GacS/GacA mutations are specific to the rhizosphere environment

In order to assess whether the observed mutations specifically conferred an advantage in the rhizosphere environment, we compared the fitness of evolved mutualists on plant roots and on liquid growth culture media. To this end, the fitness of two evolved *gacA* (Mutualists 1 and 2; ID 242 and ID 220, respectively, Table S2), and one *gacS* genotype (Mutualist 1, ID 222, Table S2) was compared relative to their direct ancestral genotypes without *gac* mutations (ID 133, ID 28 and ID 66, respectively, Table S2) within the same plant selection lines. Fitness was determined as the relative competitive fitness in direct pairwise competitions as a deviation from the initial 1:1 ancestor-to-successor ratio *in vivo* on *A. thaliana* roots and *in vitro* in Kings’ B (KB), lysogeny broth (LB), and tryptic soy broth (TSB) growth media. Post-competitive genotype ratios were determined using PCR-based high-resolution melting profile (RQ-HRM) analysis (see Methods and Materials and Supplementary figure 7). It was found that all three *gacS/gacA* mutants had a higher fitness in the rhizosphere relative to their direct ancestral genotypes without *gac* mutations (F_3, 32_ = 10.03, *P* < 0.001, Fig. 5). Interestingly, this advantage was smaller for one of the *gacA* mutants (ID 220, F_2, 6_ = 15.35, *P* = 0.004, Fig. 5) likely because its direct ancestor already showed mutualistic behaviour due to a mutation in the *accC* gene, which likely reduced the relative benefit of the *gacA* mutation within this lineage (Table S2). While the fitness benefits of *gac* mutations were mainly observed in the rhizosphere, two *gac* mutants showed improved competitive fitness in KB media indicative of general metabolic adaptations (genotype × measurement environment: F_6, 24_ = 13.02, *P* < 0.001; genotype comparisons in KB media: F_2, 6_ = 162.6, *P* < 0.001). Together, these results confirm that the genetic changes underlying the evolution of bacterial mutualism were primarily driven by plant selection.

**Figure 5.**
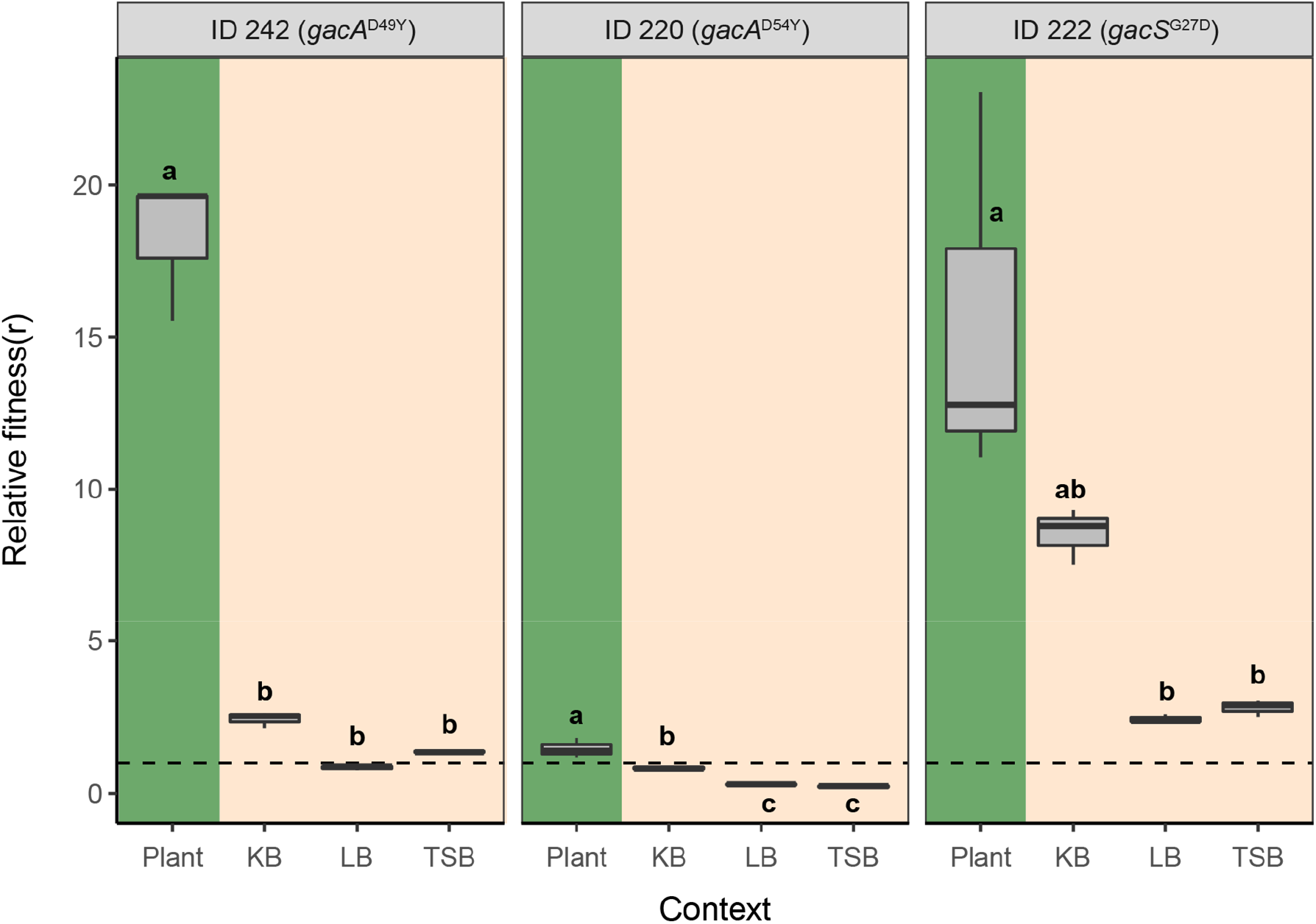
Competitive fitness of *gac* mutants relative to their direct ancestors in the rhizosphere and in *in vitro* culture media. The *gac* mutants’ relative fitness (*r*) was calculated based on the deviation from the initial 1:1 genotype ratio (dashed line) after direct competition in different environments. Fitness values above the dashed line indicate a relatively higher competitive advantage of *gac* mutants relative to their ancestral genotypes without *gac* mutations (Table S2), whereas values below the dashed line denote for decreased competitive ability of evolved *gac* mutants. In all panels, green and beige backgrounds denote competition assays conducted in the rhizosphere and in standard culture media, respectively. Different small letters above the boxplots represent significant differences in *r* between growth conditions for each mutant based on three biological replicates (one-way ANOVA, Tukey’s HSD test, α=0.05).

## Discussion

Even though beneficial plant-microbe interactions are widely documented, their evolutionary origin is less well understood. Here we studied how an initially antagonistic relationship between *P. protegens* CHA0 bacterium and its host plant, *Arabidopsis thaliana*, changed during prolonged selection over six plant growth cycles (6 months). While several studies have previously reported beneficial effects of CHA0 strain on plant growth in natural soils, it initially showed antagonism towards the plant in our experimental conditions potentially due to production of phytotoxic compounds. Crucially, this interaction rapidly evolved during the experiment leading to clear phenotypic and genetic bacterial diversification and evolution of bacterial mutualists that had relatively higher competitive advantage in the rhizosphere, and positive effect on the plant growth compared to ancestral and other evolved bacterial phenotypes.

Based on our results, we suggest the following conceptual model for the evolution of mutualism. As bacterial growth in this system was dependent on plant root exudates, it is likely that reduction in the production of exoproducts, including lytic enzymes and antimicrobial secondary metabolites (Fig. S3), by mutualists had a positive effect on plant growth and the availability of plant-derived nutrients. As many metabolites produced by *Pseudomonas spp.* are potentially phytotoxic ^20^, the observed shift from antagonism to mutualism could therefore be explained by reduced toxicity to the plant. In turn, improved plant growth likely triggered selection for mutualists that were better at competing for root exudates relative to other phenotypes, or by selectively constraining the growth of non-mutualist phenotypes via certain sanctioning mechanisms. As a support for this, ‘Mutualist 1’ phenotypes evolved better at growing on plant-derived nutrients, while ‘Mutualist 2’ phenotypes evolved to activate plant-derived scopoletin production, a compound that was relatively more harmful to non-mutualistic phenotypes. Such differences between two mutualist phenotypes were also linked with subtle differences in the predicted functional effects of observed *gacS/gacA* mutations. Together, these adaptations could have created a strong selective advantage for mutualistic phenotypes as evidenced by their relatively higher abundance in the rhizosphere at the end of the selection experiment.

At a genetic level, the evolution of mutualism could be achieved with only a few successive mutations involving mainly global regulators ^29^. This result shows an interesting parallel with recent work demonstrating that the loss of a few virulence traits can turn a pathogen into a beneficial symbiont ^7^. Evolution of mutualism was also linked with a clear phenotypic and genotypic bacterial diversification, which has previously been observed in aquatic ^39^ and soil ^40^ microcosms in response to spatial heterogeneity. Here we show that such bacterial diversification can also be driven by plant selection as evidenced by direct competition assays where competitive benefit observed in the rhizosphere was reduced or completely absent in lab media *in vitro*. Interestingly, we observed a contrasting evolutionary outcome in one of the five selection lines where ‘Stress-sensitive’ genotypes were able to become dominant alongside with ‘Ancestral-like’ genotypes potentially due to their enhanced ability to form biofilm and colonise plant roots. Interestingly, none of the phenotypes was able to reach fixation in the rhizosphere. One possibility for this is that the experiment was not long enough for the selective sweeps to drive beneficial mutations into fixation. Alternatively, it is possible that multiple phenotypes were able to coexist due to negative frequency dependent selection or because they occupied different spatial niches as seen in heterogenous soil environments ^40,41^. These hypotheses could be studied directly in the future using fluorescent microscopy and tagged strains to observe diversification and genotype fluctuations in the rhizosphere both in space and time.

In summary, our results show that in addition to recruiting beneficial bacteria from multi-species microbial communities ^8–10^, plants could also change the functioning of its associated microbiota by creating strong selection for *de novo* evolution of mutualistic bacterial genotypes. Steering bacterial evolution in the rhizosphere could thus offer plants a shortcut to improve their fitness without evolving themselves ^42–44^. Future work should focus on validating our findings in more complex microbial communities where bacterial diversification could be also affected by interactions with other microbes. In conclusion, our results call for eco-evolutionary management of plant-microbe interactions in agriculture by demonstrating that plant-associated bacteria can rapidly evolve along the parasitism-mutualism continuum within a few plant growing seasons.

## Materials and Methods

### Bacterial strain and growth conditions

We used *Pseudomonas protegens* (formerly *Pseudomonas fluorescens*) ^45^ CHA0 as a model strain, which was initially isolated from tobacco roots ^46^. The strain was chromosomally tagged with GFP and a kanamycin resistance cassette to enable specific tracking of the strain and detection of contaminations ^19^. This bacterium has the genetic potential to produce various bioactive metabolites, including the plant hormone indole-3-acetic acid (IAA), antimicrobial compounds and lytic enzymes ^47^. Prior to the experiment, bacteria were grown for 48 h on a King’s medium B ^48^ (KB) agar plate supplemented with 50 μg ml^−1^ kanamycin, a single colony was randomly picked and grown for 12 h in KB at 28 °C with agitation. The cell culture was then washed for three times in 10 mM MgSO_4_ and adjusted to 10^7^ cells ml^−1^ and used as inoculant for all plants. This inoculant was also stored at −80 °C as frozen ancestral stock, from which ‘Ancestor’ isolates were picked in later experiments.

### Host plant and growth conditions

We used *Arabidopsis thaliana* ecotype Col-0 as a model host plant. Surface-sterilized seeds were first sown in Petri dishes with agar-solidified (1.5% agar (w/v)) modified Hoagland’s medium: (KNO_3_ (3 mM), MgSO_4_ (0.5 mM), CaCl_2_ (1.5 mM), K_2_SO_4_ (1.5 mM), NaH_2_PO_4_ (1.5 mM), H_3_BO_3_ (25 μM), MnSO_4_ (1 μM), ZnSO_4_ (0.5 μM), (NH_4_)_6_Mo_7_O_24_ (0.05 μM), CuSO_4_ (0.3 μM), MES (2.5 mM) and 50 μM Fe(III)EDTA, pH = 5.8) and stratified for 2 days at 4 °C after Petri dishes were positioned vertically and transferred to a growth chamber (20 °C, 10 h light/14 h dark, light intensity 100 μmol m^−2^ sec^−1^). After two weeks of incubation, two seedlings were transferred to closed and sterile ECO2 boxes (http://www.eco2box.com/ov80xxl_nl.htm) for selection experiment. The ECO2 boxes were filled with 260 g of dry, carbon-free silver sand that was previously washed with MilliQ water to remove dissolvable chemical elements and heated to 550 °C for 24 h to remove remaining organic material. Prior to transplantation the sand was amended with 13 ml of modified Hoagland medium.

### Design of the selection experiment

The selection experiment was conducted in a gnotobiotic system to remove confounding effects that may emerge as a result of competitive interactions with other micro-organisms, and to place the focus on plant-mediated selective pressures. Moreover, we allowed only the bacteria to evolve during the experiment and used new clonal plants at every bacterial transfer. We set up five independent plant-bacterium replicate lines, which were grown for six independent plant growth cycles (see Figure S1 for an overview of the experimental design). The experiment was started by inoculating 10^6^ cells of the stock *P. protegens* CHA0 culture (From here on abbreviated as “ancestor”) into the rhizosphere of two-week-old *A. thaliana* seedlings growing in sterile silver sand within ECO2 boxes (two plants per replicate selection line). Inoculated plants were then grown for four weeks (20 °C, 10 h light/14 h dark, light intensity 100 μmol m^−2^ sec^−1^) after which the plant growth cycle was terminated and root-associated bacteria were harvested by placing the roots of both plants into a 1.5 ml Eppendorf tubes filled with 1 ml 10 mM MgSO_4_ and two glass beads. Rhizosphere bacteria were suspended into the liquid using TissueLyser II at a frequency of 20 s^−1^ for 1 min after which bacterial cell densities were determined using flow cytometry (BD Accuri™ C6 Plus, thresholds for FSC: 2000, SSC: 8000). After this, 10^6^ cells were inoculated to the rhizosphere of new *A. thaliana* plants to initiate the next plant growth cycle. Possible contaminations were checked by plating the suspension on 3 g l^−1^ tryptic soy agar (TSA) plates and it was verified that all colonies carried the *GFP* marker gene, as observed under UV light.

### Bacterial life-history traits measurements

Individual bacterial colonies were isolated from all replicate plant selection lines for life-history measurements at the end of the second, fourth and sixth plant growth cycle by dilution plating the rhizosphere suspension on 3 g l^−1^ TSA plates. After incubation at 28 °C for 24 h, 16 colonies were randomly picked from each replicate selection lines resulting in a total of 240 evolved and 16 ancestral colonies. All these colonies were characterized for a set of key bacterial life-history traits representative of growth, stress resistance and traits linked with plant-microbe interactions.

#### a. Bacterial growth yield in KB medium

All the bacterial isolates were grown in 96-well plates with 160 μl 1/3 strength liquid KB, at 20 °C without shaking. Bacterial yield was determined as the maximum optical density at 600 nm after three days of growth using a spectrophotometer (SPECTROstar Nano).

#### b. Bacterial stress resistance

We measured bacterial resistance to a range of different stresses using various 96-well microplate assays. Abiotic stress resistance was determined by growing bacteria in 160 μl of 1 g l^−1^ tryptic soy broth (TSB) containing 0.0025% H_2_O_2_ (oxidative stress), 15% polyethylene glycol (PEG)-6000 (water potential stress) or 2% NaCl (salt stress). We used resistance to antibiotics commonly produced by rhizosphere microorganisms as indicator of biotic stress resistance. Antibiotic resistance was tested in 160 μl of 1 g l^−1^ TSB supplemented with 1 μg ml^−1^ streptomycin, 1 μg ml^−1^ tetracycline, or 5 μg ml^−1^ penicillin, respectively. Bacterial growth with and without stresses were determined after three days of growth at 20 °C without shaking as optical density at 600 nm and stress resistance defined as the ratio of bacterial growth in the stressed relative to the non-stressed control treatment.

#### c. Traits linked with plant-microbe interactions

*P. protegens* CHA0 harbours several traits that are linked to plant growth including production of antibiotics and plant hormones. To assess these traits, we grew each bacterial colony in 96-well plates containing 160 μl of 1/3 strength liquid KB per well at 20 °C with agitation for 72 h. Cell-free supernatants were obtained by filter sterilization (0.22 μm) using Multiscreen HTS 96-well filtration plates (1000 × g, 30 min), which were used to measure the production of the plant hormone auxin (Indole-3-acetic acid (IAA)), iron-chelating siderophores and proteolytic activity. Furthermore, we also measured antifungal and antibacterial activity of all colonies.

##### IAA detection

The production of the plant hormone auxin was determined with a colorimetric test ^49^. Briefly, 30 μl *P. protegens* CHA0 cell-free filtrate was incubated with 30 μl R1 reagent (12 g l^−1^ FeCl_3_, 7.9 M H_2_SO_4_) for 12 h in the dark and optical density read at 530 nm of the colorimetric complex was used as a measurement of IAA concentration.

##### Siderophore activity

Iron chelating ability was measured as a proxy for siderophore production ^50^. To this end, 100 μl of *P. protegens* CHA0 cell-free filtrate was mixed with 100 μl of modified CAS solution (with 0.15 mM FeCl_3_) and optical density read at 630 nm after 3 hours of incubation was used as a proxy of siderophore production. The iron chelating ability was calculated based on the standard curve based on modified CAS assay solution with a range of iron concentration (0, 0.0015, 0.003, 0.006, 0.009, 0.012, 0.015 mM FeCl_3_).

##### Proteolytic activity

The proteolytic activity assay we used was adapted from Smeltzer et al. (1993) ^51^. Briefly, 15 μl of *P. protegens* CHA0 cell-free filtrate was incubated with 25 μl of azocasein (2% w/v in 50 mM Tris-HCl pH 8.0) at 40 °C for 24 hours. 125 μl of 10% w/v cold trichloroacetic acid (TCA) was added to precipitate superfluous azocasein, and then 100 μl supernatant was neutralized with 100 μl of 1M NaOH after centrifugation at 5000 rpm for 30 minutes. Optical density read at 440 nm was used as a proxy of exoprotease activity.

##### Tryptophan side chain oxidase (TSO) activity

TSO activity, an indicator of quorum sensing activity in *P. protegens* CHA0, was measured based on an modified established colorimetric assay ^52^: Three-day-old bacterial cultures grown in 1/3 strength liquid KB were mixed at a 1:1 ratio with a reagent solution (5 g l^−1^ SDS, 37.6 g l^−1^ glycine 2.04 l^−1^ g tryptophan, pH 3.0) and TSO activity was measured as optical density at 600 nm after overnight incubation.

##### Biofilm formation

We quantified bacterial biofilm formation using a standard protocol ^53^. Briefly, bacteria were grown at 20 °C for 72h in 160 μl 1 g l^−1^ TSB in 96-well microtiter plate with TSP lid (TSP, NUNC, Roskilde, Denmark). Planktonic cells were removed by immersing the lid with pegs three times in phosphate-buffered saline solution (PBS). Subsequently, the biofilm on the pegs was stained for 20 minutes in 160 μl 1% Crystal Violet solution. Pegs were washed five times in PBS after which the Crystal Violet was extracted for 20 minutes from the biofilm in a new 96-well microtiter plate containing 200 μl 96% ethanol per well. Biofilm formation was defined as the optical density at 590 nm of the ethanol extracted Chrystal Violet ^54^.

##### Inhibition of other microorganisms

Antimicrobial activity was defined as the relative growth of the target organism in *P. protegens* supernatant compared to the control treatment. Antifungal activity of the cell-free supernatant was assessed against the ascomycete *Verticillium dahlia*. The fungus was grown on potato dextrose agar (PDA) at 28 °C for 4 days, after which plugs of fungal mycelium were incubated in potato dextrose broth (PDB) medium at 28 °C and gentle shaking for 5 days. Fungal spores were collected by filtering out the mycelium from this culture over glass wool. Subsequently, spores were washed and resuspended in water and the OD_595_ of the suspension was adjusted to 1. Five μl of this spore suspension was then inoculated with 15 μl *P. protegens* CHA0 cell-free filtrate and incubated in 160 μl of 1g l^−1^ PDB medium for 2 days at 20 °C in 96 well plates. Fungal growth was measured as optical density at 595 nm after 2 days of growth and contrasted with the growth in the control treatment (PDB medium without *P. protegens* supernatant). Antibacterial activity was determined using the plant pathogen *Ralstonia solanacearum* as a target organism. *R. solanacearum* was grown in 160 μl of 1 g l^−1^ TSB medium supplemented with 15 μl of *P. protegens* CHA0 cell-free filtrate or 15 μl of 1/3 strength liquid KB as a control for 2 days at 20 °C. *R. solanacearum* growth was measured as optical density at 600 nm.

### Determining changes in *P. protegens* CHA0 interactions with *A. thaliana* after the selection experiment

Based on the life-history trait measurements, five distinct bacterial phenotypes were identified using K-means clustering analysis (Fig S2). In order to assess whether phenotypic changes reflected shifts in the strength and type of plant-bacterium interaction, we chose five isolates from each bacterial phenotype group representing each replicate selection line and five ancestral isolates for further measurements (a total of 30 isolates, Table S2).

#### a. Effects of ancestor and evolved bacteria on plant performance

For each isolate we measured root colonizing ability and impact on plant performance. All 30 bacterial isolates were incubated overnight in 1/3 KB strength liquid at 20 °C. The culture was centrifuged twice for five minutes at 5000 x g and the pellet was washed and finally resuspended in 10 mM MgSO_4_. The resulting suspension was adjusted to an OD_600_ of 0.01 for each strain as described previously ^55^. Ten μl of the bacterial suspension (or 10 mM MgSO_4_ as a control) was applied to the roots of three 10-day old sterile *Arabidopsis thaliana* Col-0 seedlings (excluding 2-days of stratification at 4 °C) grown on vertically positioned Petri dishes with agar-solidified (1.5% agar (w/v)) modified Hoagland’s medium (n = 3 biological plant replicates, each containing 3 seedlings). Plants were grown for 14 days before harvesting. Plants were photographed before and 14 days after bacterial inoculation.

Bacterial effects on plant health were quantified as leaf ‘greenness’ as the presence of ancestral strain was observed to lead to bleaching and loss of chlorophyll in *A. thaliana* leaves. The ‘greenness’ was quantified from photographs by measuring the number of green pixels. To this end, photographs were first transformed in batch using Adobe Photoshop 2021 by sequentially selecting only green areas followed by thresholding balancing green tissue over background noise (Level 80). This resulted in black-and-white images for further analysis, and the mean number of white pixels per fixed-sized region-of-interest of the aboveground tissue was subsequently determined as ‘greenness’ using ImageJ. The numbers of lateral roots and the primary root length were also measured using ImageJ (version 1.50i). The root morphology data measured at the end of the experiment was normalized with the data collected at the time of inoculation for each individual seedling.

To determine shoot biomass, the rosette of each plant was separated from the root system with a razor blade and weighted. The roots were placed into a pre-weighted 1.5 ml Eppendorf tubes to quantify the root biomass. Then these tubes were filled with 1 ml 10 mM MgSO_4_ buffer solution and two glass beads. The rhizosphere bacteria were suspended into buffer solution using TissueLyser II at a frequency of 20 s^−1^ for 1 min after which bacterial densities were determined using flow cytometry as described above.

Shoot biomass, root biomass, root length, and number of lateral roots were used in a principal component analysis (PCA) to calculate an overall impact of the bacteria on plant performance (Fig 2E). The first principal component (PC1) explained 79.9% of the variation and was normalized against the control treatment to be used as a proxy of ‘Plant performance’ in which positive values reflect plant growth promotion and negative values plant growth inhibition.

#### b. Root derived carbon source utilization

To measure changes in bacterial growth on potential root derived carbon sources, we measured the growth of all 256 isolates using modified Ornston and Stanier (OS) minimal medium ^56^ supplemented with single carbon sources at a final concentration of 0.5g l^−1^ in 96-well plates containing 160 μl carbon supplemented OS medium per well. The following carbon sources were selected based on their relatively high abundance in *Arabidopsis* root exudates^21^: alanine, arabinose, butyrolactam, fructose, galactose, glucose, glycerol, glycine, lactic acid, putrescine, serine, succinic acid, threonine and valine. Bacterial growth was determined by measuring optical density at 600 nm after three days incubation at 20 °C.

#### c. GUS *histochemical staining assay and bacterial growth under scopoletin stress*

To investigate effects of the ancestor and evolved strains of *P. protegens* CHA0 on expression of *MYB72* gene, we applied *GUS* histochemical staining assay to the 30 selected isolates (Table S2). MYB72 is a transcription factor involved in production of the coumarin scopoletin in *Arabidopsis* roots and specific rhizobacteria can upregulate expression of *MYB72* in the roots. Scopoletin is an iron-mobilizing phenolic compound with selective antimicrobial activity ^22^. Seedlings of the *A. thaliana MYB72_pro_:GFP-GUS* ^24^ reporter line were prepared as described above. Seven day old seedlings were inoculated directly below the hypocotyls with 10 μl of a bacterial suspension (OD660 = 0.1) as described previously ^24^. At 2 days after inoculation, the roots were separated from the shoots and washed in Milli-Q water (Milliport Corp., Bedford, MA) to remove all the adhered bacteria. GUS staining of the roots was performed in 12-well microtiter plates where each well contained roots of 5 to 6 seedlings and 1 mL of freshly prepared GUS substrate solution (50 mM sodium phosphate with a pH at 7, 10 mM EDTA, 0.5 mM K4[Fe(CN)6], 0.5 mM K3[Fe(CN)6], 0.5 mM X-Gluc, and 0.01% Silwet L-77) as described previously ^57^. Plates were incubated in the dark at room temperature for 16 h. The roots were fixed overnight in 1mL ethanol:acetic acid (3:1 v/v) solution at 4 ℃ and transferred to 75% ethanol. Then the pictures of each microtiter plates were taken, and GUS activity was quantified by counting the number of blue pixels in each well of the microtiter plates using image analysis in ImageJ (version 1.52t). To assess the effects of scopoletin on ancestral and evolved *P. protegens* CHA0 isolates, we applied a sensitivity assay to the 30 selected isolates (Table S2). In brief, growth of bacterial isolates was measured in 1 g l^−1^ TSB medium (160 μl) supplemented with scopoletin at final concentrations of 0 μM (control), 500 μM, 1000 μM, and 2 mM using optical density at 600 nm after 96 h incubation at 20 °C without shaking in 96-well microtiter plates. Maximal effect (Emax) of scopoletin was calculated via R package ‘GRmetrics’ ^58^ as an indication of scopoletin tolerance.

### Whole genome sequencing

All 30 isolated phenotypes were whole genome sequenced to identify possible mutations and affected genes. To this end, isolates were cultured overnight at 28 °C in 1/3 strength liquid KB. Chromosomal DNA was isolated from each culture using the GenElute™ Bacterial Genomic DNA Kit Protocol (NA2100). DNA samples were sheared on a Covaris E-220 Focused-ultrasonicator and sheared DNA was then used to prepare Illumina sequencing libraries with the NEBNext® Ultra™ DNA Library Prep Kit (New England Biolabs. France) and the NEBNext® Multiplex Oligos for Illumina® (96 Index Primers). The final libraries were sequenced in multiplex on the NextSeq 500 platform (2 x 75 bp paired-end) by the Utrecht Sequencing Facility (http://www.useq.nl) yielding between 1.0 and 6.4 million reads per sample equivalent to ~10 – 70 fold coverage (based on comparison with the original 6.8 Mbp reference genome NCBI GenBank: <u>CP003190.1</u>).

### Variant calling analysis

We first constructed an updated reference genome of *P. protegens* CHA0, carrying the *GFP* marker gene on its chromosome, from the ancestral strain using the A5 pipeline with default parameters ^59^. The input dataset for this sample consisted of 3,1 M reads and totals an approximate 34-fold coverage. The size of the updated reference genome is 6.8 Mbp, with a G+C content of 63.4%, and it comprises 80 scaffolds, with a *N*_50_ value of 343 Kbp. We subsequently used PROKKA ^60^ (version 1.12; https://github.com/tseemann/prokka) for full annotation of the updated reference genome, and this resulted in the identification of 6,147 genes. The updated genome is deposited in NCBI GenBank with following reference: <u>RCSR00000000.1</u>.

Having established the ancestral genome sequence, we subsequently used Snippy (version 3.2-dev; https://github.com/tseemann/snippy) to identify and functionally annotate single nucleotide polymorphisms and small insertions and deletions (indels) for each individual strain. In addition, we investigated the breadth of coverage for each gene per sample with BedTools ^61^ to identify genes with large insertions or deletions. An overview of the polymorphisms is shown in Supplementary Table S3. Raw sequencing data for this study is deposited at the NCBI database under BioProject <u>PRJNA473919</u>.

### Relative competitive fitness of *gac* mutants measured *in vivo* and *in vitro*

The relative competitive fitness of selected *gac* mutants was measured in direct competition with their direct ancestors both *in vivo* in the rhizosphere of *A. thaliana* and *in vitro* in different standard culture media. Relative fitness was measured as deviation from initial 1:1 ratio of bacterial clone pairs based on PCR-based high-resolution melting profile (RQ-HRM) analysis. Three pairs of isolates were selected: A) Evolved *gacA* ID 242 (genotype *oafA*^Y335X^ · *RS17350*^A77A.fsX14^ · *gacA*^D49Y^) and its direct ancestral genotype 133 (genotype *oafA*^Y335X^ · *RS17350*^A77A.fsX14^) from evolutionary line 1; B) Evolved *gacA* ID 220 (genotype *galE*^V32M^ · *accC*^E413K^ · *gacA*^D54Y^) and its direct ancestral genotype 28 (genotype *galE*^V32M^ · *accC*^E413K^) from line 2; C) Evolved *gacS* ID 222 (genotype *oafA*^K338S.fsX18^ · *gacS*^G27D^) and its direct ancestral genotype 66 (genotype *oafA*^K338S.fsX18^) from line 3. Bacterial isolates were first grown overnight in KB medium at 28 °C, centrifuged at 4,500 rpm for 10 min and the pellet resuspended in 10 mM MgSO_4_. This washing procedure was repeated twice. The resulting bacterial suspensions were diluted to OD_600_ = 0.05. The initial inoculum for the competition assays was then generated by mixing equal volumes of evolved and ancestral competitors in a ratio of 1:1.

#### Measuring competitive fitness in A. thaliana rhizosphere

This assay was performed on the roots of 10-day old *A. thaliana* seedlings grown on full strength Hoagland agar plates, which were prepared as described earlier. Twenty μl of the initial inoculum, containing a total of 10^6^ bacterial cells, was inoculated on to the root-shoot junction of each seedling. After 14 days of growth, bacterial populations were isolated from the roots as described earlier and stored at −80 °C in 42.5% glycerol for relative abundance measurements.

#### Measuring competitive fitness in culture media

Competition assays were also performed in three commonly used nutrient-rich growth media: Kings’ B (KB), lysogeny broth (LB), and tryptic soy broth (TSB). KB contained 20 g proteose peptone, 1.5 g MgSO_4_.7H_2_O, 1.2 g KH_2_PO_4_ and 10 g glycerol per litre and the pH was adjusted to 7.3 ± 0.2. TSB contained 30 g tryptic soy broth per litre and pH was adjusted to 7.3 ± 0.2. LB contained 10 g peptone, 5 g yeast extract and 5 g NaCl per litre. Twenty μl inoculum of competing strains, containing about 10^6^ bacterial cells, were added into wells containing 140 μl fresh medium in a 96-well plate. The microplates were incubated at 28 °C without shaking for 48 after 80 μl sample was harvested and stored at −80 °C in 42.5% glycerol from each well for relative abundance measurements.

#### RQ-HRM assay for quantifying changes in genotype frequencies after competition

We used a High-Resolution Melting (HRM) curve profile assay with integrated LunaProbes to quantify the ratio of mutant to wild type genotypes ^62–64^. The probes and primers used in this study are listed in Table S4. Primers were designed using Primer3. Probes were designed with the single nucleotide polymorphism (SNP) located in the middle of the sequence, and the 3’ end was blocked by carbon spacer C3. The primer asymmetry was set to 2:1 (excess primer: limiting primer) in all cases. Pre-PCR was performed in a 10-μl reaction system, with 0.25 μM excess primer, 0.125 μM limiting primer, 0.25 μM probe, 0.5 μl bacterial sample culture (100-fold diluted saved sample, OD_600_ is about 0.01), 1X LightScanner Master Mix (BioFire Defense). DMSO with the final concentration 5% was supplemented in all reactions to ensure the targeted melting domains are within the detection limit of the LightScanner (Idaho Technology Inc.). Finally, MQ water was used to supplement up to 10 μl. A 96-well black microtiter plate with white wells was used to minimize background fluorescence. Before amplification, 25 μl mineral oil was loaded in each well to prevent evaporation, and the plate was covered with a foil seal to prevent the degradation of fluorescent molecules. Amplification was initiated by a holding at 95 °C for 3 min, followed by 55 cycles of denaturation at 95 °C for 30 s, annealing at 60 °C for 30 s and extension at 72 °C for 30 s and then kept at 72 °C for 10 min. After amplification, samples were heated in a ThermalCycler (Bio-Rad) shortly to 95 °C for 30 s to denature all double-stranded structures followed by a rapid cooling to 25 °C for 30 s to facilitate successful hybridization between probes and the target strands. The plate was then transferred to a LightScanner (Idaho Technology Inc.). Melting profiles of each well were collected by monitoring the continuous loss of fluorescence with a steady increase of the temperature from 35 °C to 97 °C with a ramp rate of 0.1 °C /s. The relative quantification was based on the negative first derivative plots using software MATLAB. The areas of probe-target duplexes melting peaks were auto-calculated by ‘AutoFit Peaks I Residuals’ function in software PeakFit (SeaSolve Software Inc.). The mutant frequency *X* was calculated using the formula shown below:

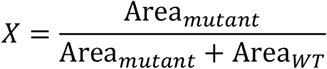

To validate the RQ-HRM method, standard curves were generated by measuring mixed samples with known proportions of mutant templates: 0%, 10%, 20%, 30%, 40%, 50%, 60%, 70%, 80%, 90% and 100%. Measurements for each sample were done in triplicate. Linear regression formula of each mutant between actual frequencies and measured frequencies were shown in Figure S7. The high R^2^ values, and nearly equal to 1 slope values of these equations, confirmed that the RQ-HRM method can accurately detect mutants’ frequency in a mixed population.

The relative fitness of the evolved strains was calculated according to previous studies using the following equation ^65,66^:

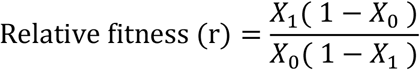

*X*_0_: initial mutant frequency; (1−*X*_0_): initial ancestor frequency. *X*_1_: final mutant frequency; (1−*X*_1_): final ancestor frequency.

## Acknowledgments

We thank Roy van der Meijs for his excellent work on the scopoletin sensitivity assays and Ke Yu, Roeland Berendsen and members of the Plant-Microbe Interactions lab for helpful discussion. This work was supported by a China Scholarship Council fellowship (to E.L.), a postdoctoral fellowship of the Research Foundation Flanders (FWO 12B8116RN) (to R.D.J.) and Royal Society Research Grants (grant nos. RSG\R1\180213 and CHL\R1\180031) at the University of York (V-P.F).

## Author contributions

EL, PB and AJ designed the experiments. EL, HJ and CL performed the experiment. EL, RJ and AJ analysed the data. All authors collegially wrote the manuscript.

## Declaration of Interests

Authors declare no competing interests.

## Supplementary Materials

### Data and materials availability

The *P. protegens* CHA0-GFP reference strain genome sequence, determined for this study, is deposited on GenBank: RCSR00000000.1. Raw sequencing data used in this study are deposited at the NCBI database under BioProject PRJNA473919. Raw data of *P. protegens* CHA0 phenotypic traits, Supplementary dataset 1 and 2, are deposited at Mendeley Data: DOI: 10.17632/wh3ytm5rn8.1

### Supplementary figures

**Figure S1.**
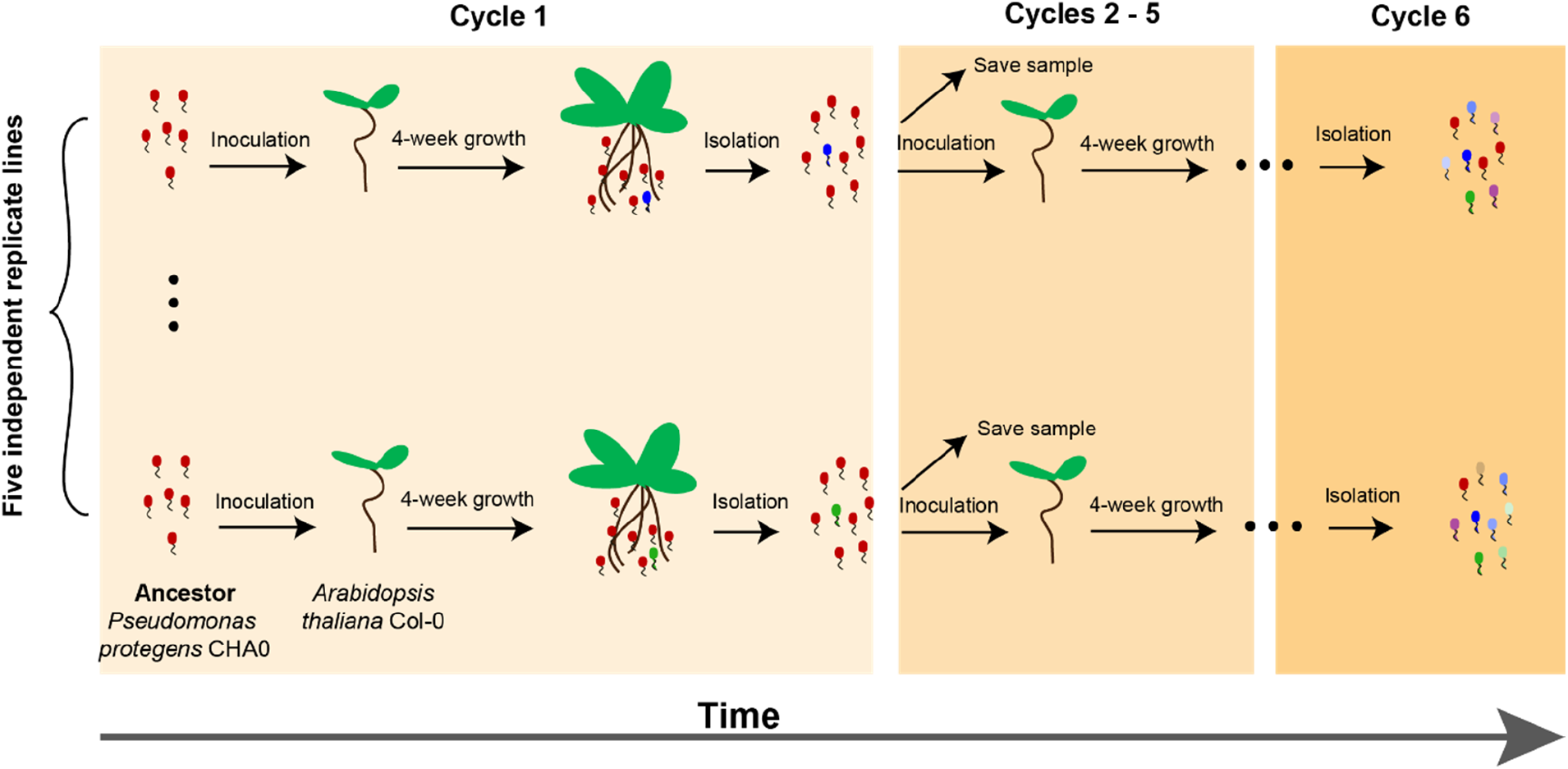
Overview of the experimental design. In this study, we experimentally evolved *Pseudomonas protegens* CHA0 in the rhizosphere of sterile *Arabidopsis thaliana* plants. We used a gnotobiotic, organic carbon-free soil system in which bacterial fitness strictly depended on their interaction with plants. We set up five independent plant replicate lines, which were passed over six plant growth cycles (4 weeks each). To this end, 10^6^ cells of the ancestral *P. protegens* CHA0 strain were introduced to the rhizosphere of two *A. thaliana* seedlings grown in sterile silver sand supplemented with a plant nutrient solution in sterile ECO2 boxes. At the end of each growth cycle, the rhizosphere bacterial population was harvested, and 10^6^ cells were inoculated onto a new plant. The remaining bacteria were kept as frozen stock at −80 °C. At the end of the experiment, bacteria from the −80 °C stock were plated on 3 g l^−1^ Tryptic Soy Agar and sixteen single bacterial isolates were randomly selected from each replicate line at the end of the second, fourth and sixth growth cycle. In total, 240 evolved isolates and sixteen ancestral isolates were phenotyped regarding traits associated with bacterial fitness and mutualistic activity with the plant.

**Figure S2.**
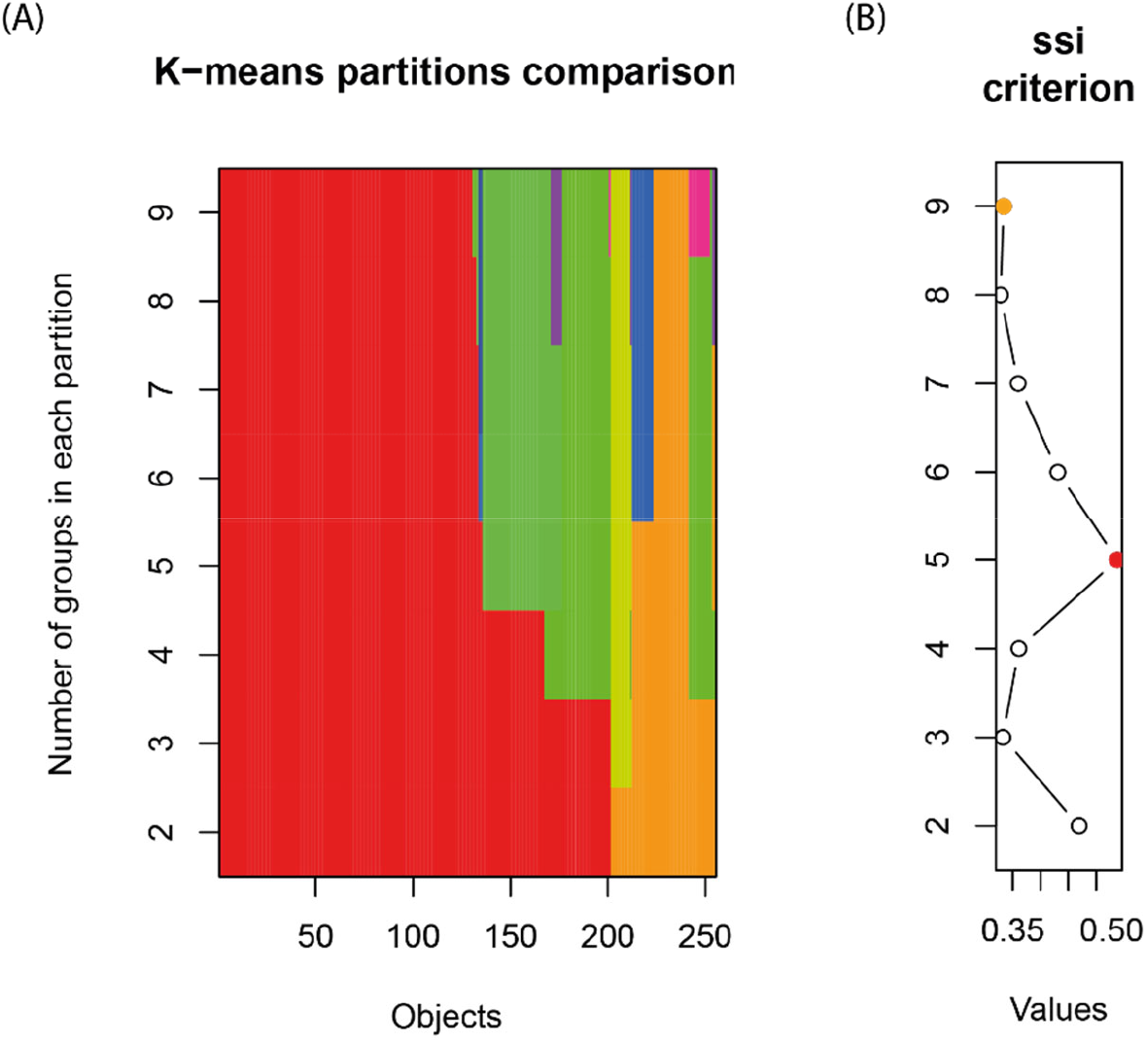
K-means clustering analysis of evolved *Pseudomonas protegens* CHA0 isolates. Isolated colonies were classified based on 14 phenotypic traits associated with bacterial fitness in the rhizosphere and mutualistic activity with the plant. In panel A, the x-axis (“Objects”) represents the 256 screened isolates while the y-axis represents the potential number of clusters (K) shown in different colours. Panel B shows the SSI criterion values indicating the most parsimonious number of clusters needed to classify isolates into distinct phenotypic groups. Based on this analysis, we classified the isolates into five clusters (the highest SSI criterion value).

**Figure S3.**
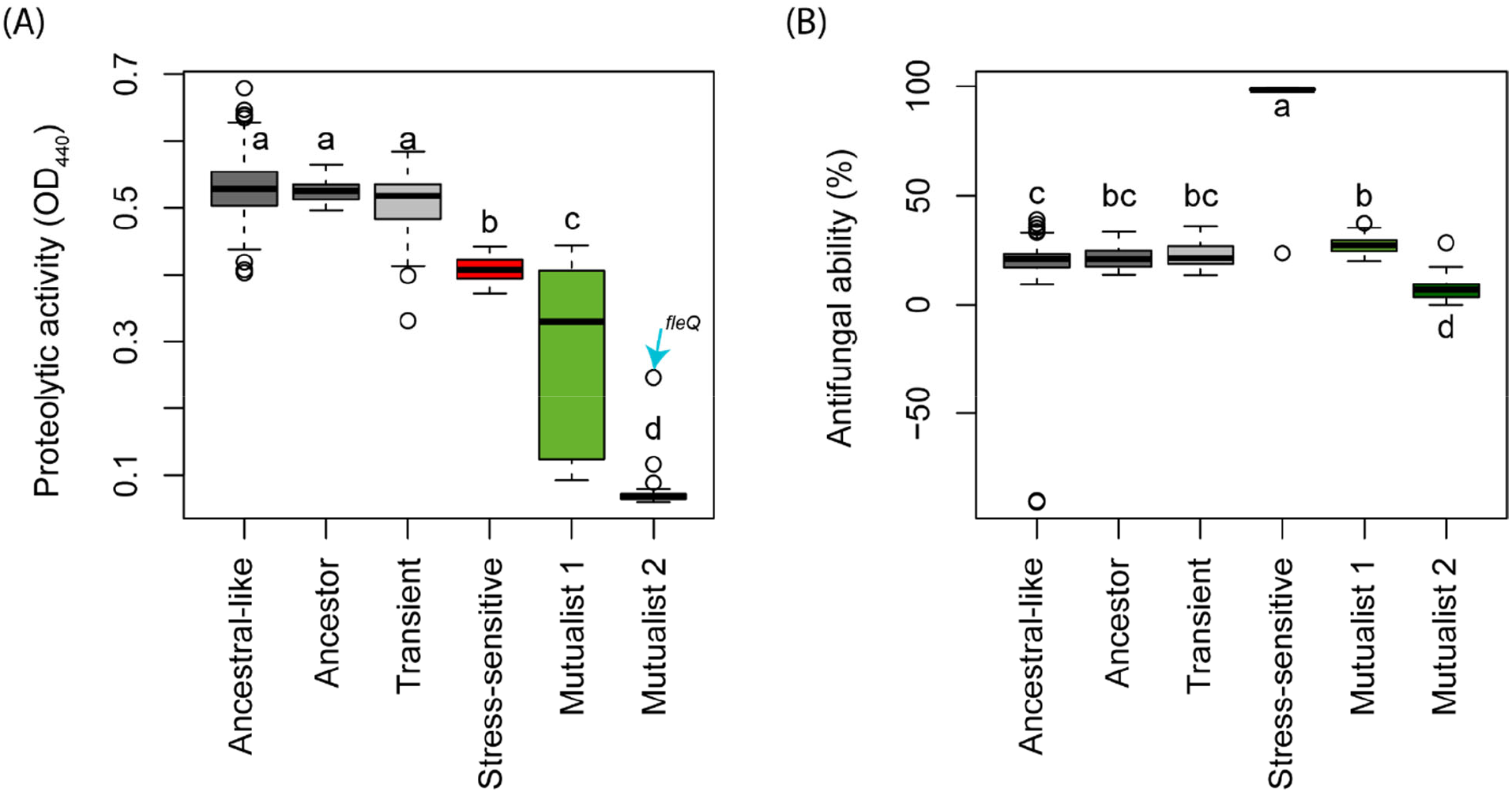
Comparing the differences in extracellular proteolytic and antifungal activity of ancestral and evolved *Pseudomonas protegens* CHA0 isolates. We measured the proteolytic (A) and antifungal activity (B) as a proxy for secondary metabolite production. In total, we characterized 16 ancestral, 119 ancestral-like, 11 stress-sensitive, 37 mutualist 1, 31 mutualist 2 and 41 transient isolates (Supplementary dataset 1). In panel A, the blue arrow indicates one isolate that was phenotypically clustered as mutualist 2, but genetically bearing a unique *fleQ* mutation. Statistical testing was carried out using ANOVA. Different letters indicate significant differences based on a Tukey HSD test (α=0.05).

**Figure S4.**
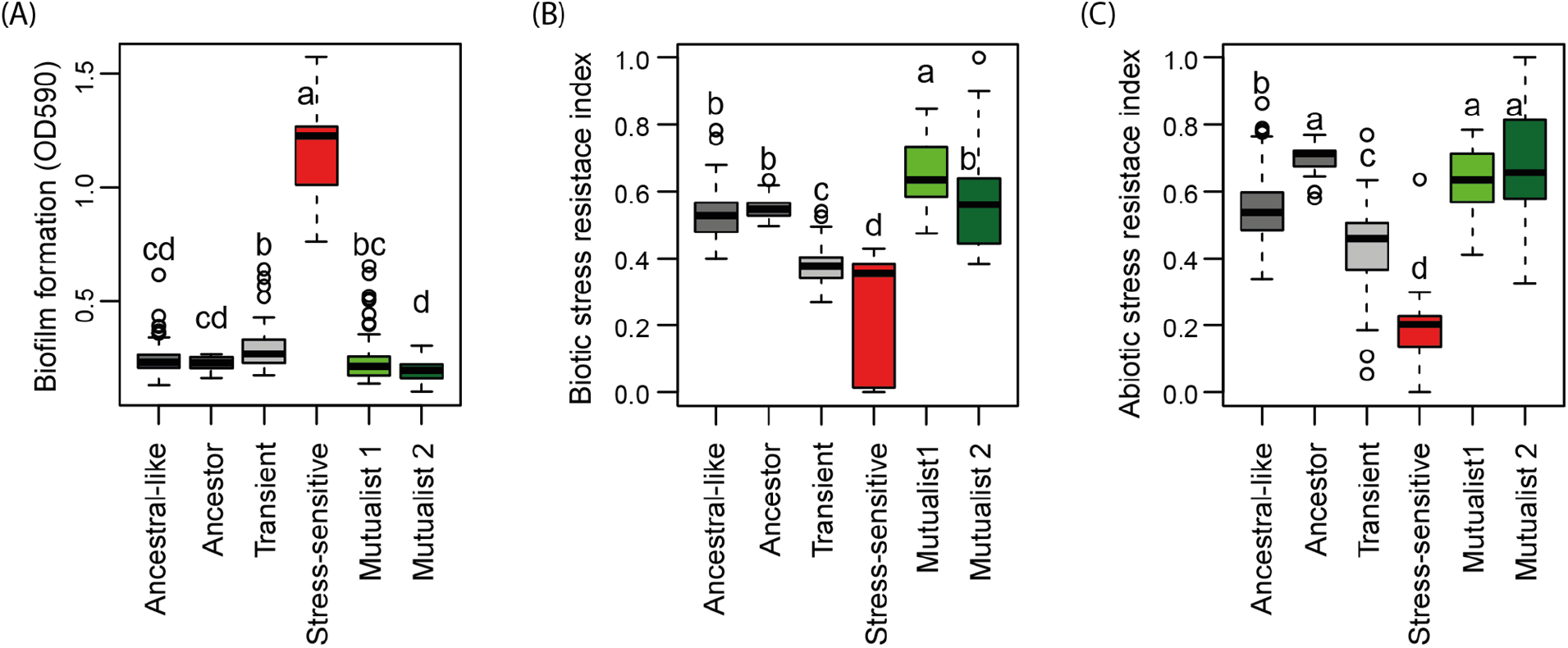
Comparing biofilm formation, biotic and abiotic stress resistance of ancestral and evolved *Pseudomonas protegens* CHA0 isolates. In total, we characterized 16 ancestral, 119 ancestral-like, 11 stress-sensitive, 37 mutualist 1, 31 mutualist 2 and 41 transient isolates (Supplementary dataset 1). Panels A to C show biofilm formation, biotic stress resistance index (normalised PC1 of combined ability to grow in the presence of sub lethal doses of the antibiotics streptomycin, tetracycline and penicillin) and abiotic stress resistance index (normalised first principal component of combined ability of each isolate to grow under oxidative stress, water potential stress and salt stress) respectively. Statistical testing was carried out using ANOVA. Different letters indicate significant differences based on a Tukey HSD test (α=0.05).

**Figure S5.**
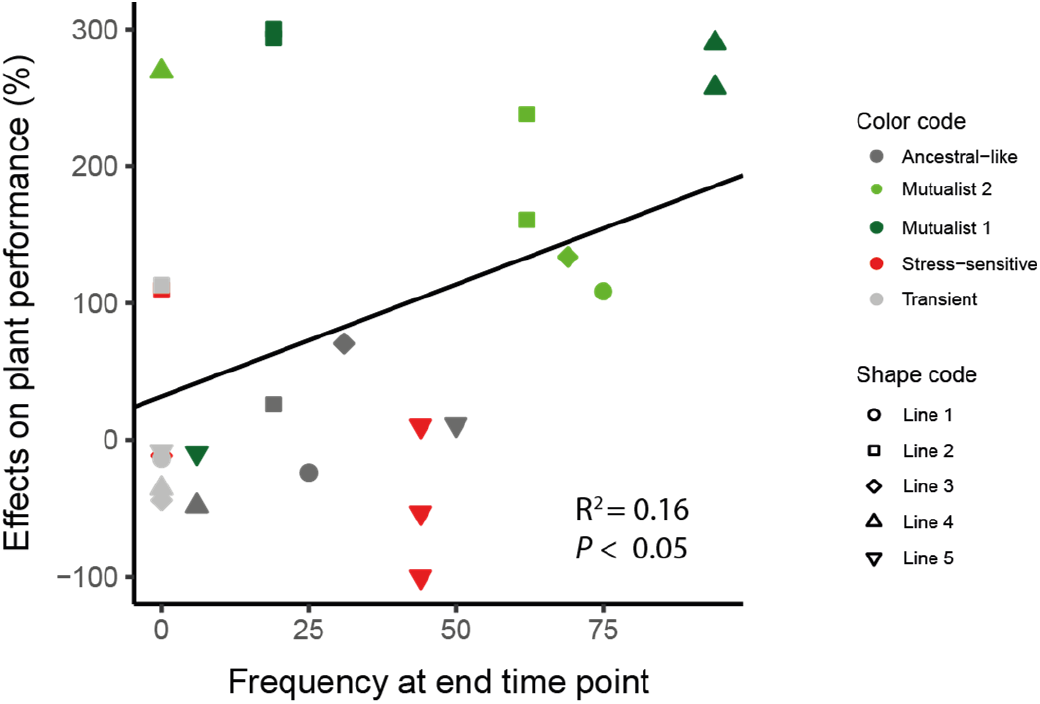
Positive relationship between phenotype frequency at the end of the selection experiment and isolate effect on the plant performance. Five representative bacterial isolates from each phenotype in addition to the ancestor were selected to measure their effects on *Arabidopsis thaliana* growth in terms of combined ‘Plant performance’ index (30 isolates altogether, each replicate line represented; See Table S2). The y-axis represents the beneficial effect of isolates on plant performance relative to the ancestor. Values on the x-axis show the relative abundance of evolved phenotypes in their respective selection lines at the end of the sixth plant growth cycle (see Figure 1). Colours and shapes represent different phenotypes and selection lines, respectively.

**Figure S6.**
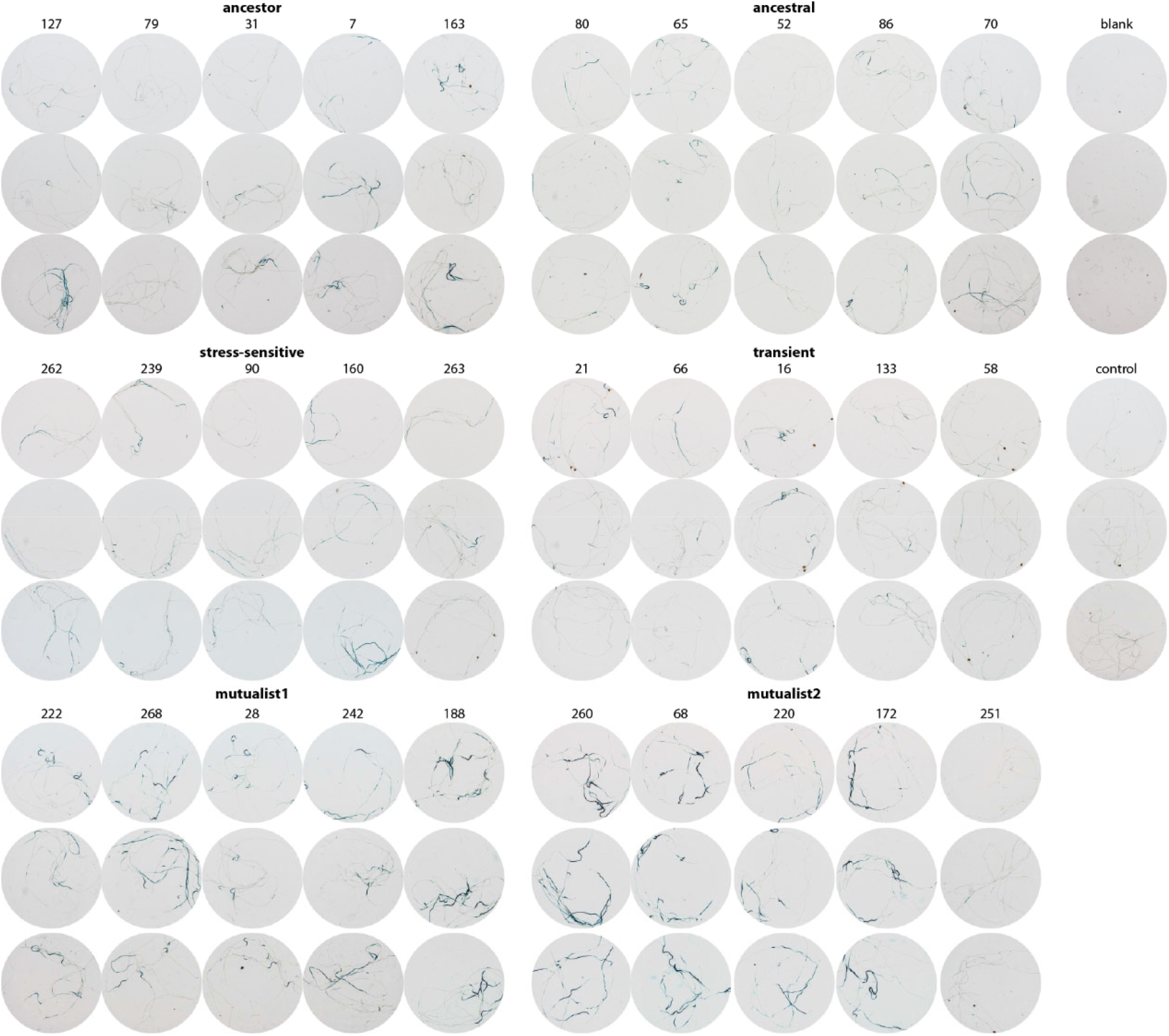
Induction of *MYB72* assayed in a GUS reporter line in *Arabidopsis thaliana* by the ancestor and a subset of evolved *Pseudomonas protegens* CHA0 isolates. GUS staining was performed at 2 days after bacterial inoculation (n = 3 biological plant replicates, each containing 5 to 6 seedlings). See Table S2 for detailed information of the isolates.

**Figure S7.**
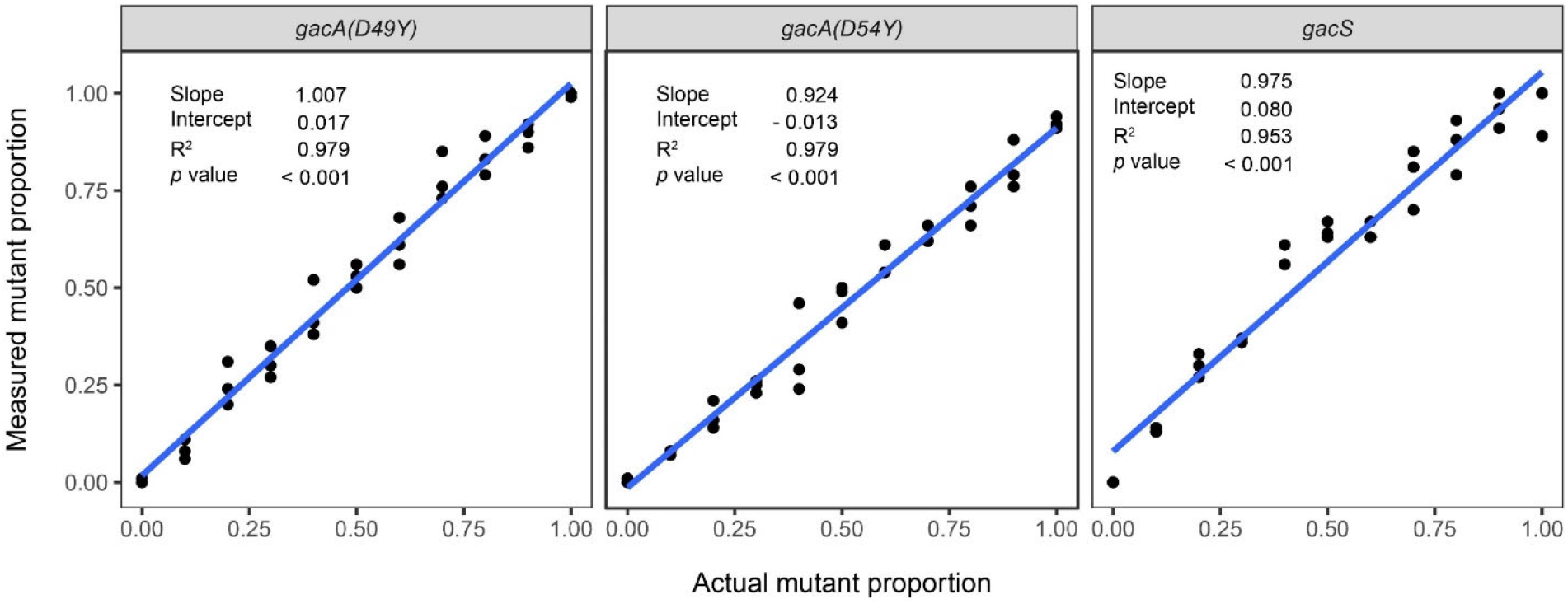
Standard curves of measured mutant versus ancestor proportion as a function of the actual proportion, using series of mixed samples with known proportions (0%, 10%, 20%, 30%, 40%, 50%, 60%, 70%, 80%, 90% and 100% of mutant templates). Relative densities of mutant *gacA*^D49Y^, *gacA*^D54Y^, and *gacS*^G27D^ were measured by PCR-based high-resolution melting profile (RQ-HRM) analysis. Measurements for each sample were done in triplicate. In each plot, the black dots represent the measurements, the blue fit line was generated based on linear regression model.

**Table S1.** *In vitro* measurement of different aspects of bacterial life-history traits, including bacterial growth, tolerance to diverse abiotic and biotic stresses, production (or activity) of bioactive compounds and antimicrobial activity. In total, 14 different phenotypic traits were measured for 256 *Pseudomonas protegens* CHA0 isolates in this study including 16 isolated isolates of ancestor.

| Aspects of life-history traits | Details of measured traits |
| --- | --- |
| Bacterial growth yield | Bacterial growth yield in King's medium B |
| Stress tolerance | Growth yield under abiotic stresses: oxidative stress, water potential stress, salt stress<br>Growth yield under biotic stresses (antibiotics): streptomycin stress, tetracycline stress, penicillin stress |
| Production (or activity) of bioactive compounds | IAA (auxin) production, siderophore activity, quorum sensing (TSO) activity, proteolytic activity, biofilm formation |
| Antimicrobial activity | Antifungal activity: *Verticillium dahliae*<br>Antibacterial activity: *Ralstonia solanacearum* |

**Table S2.**
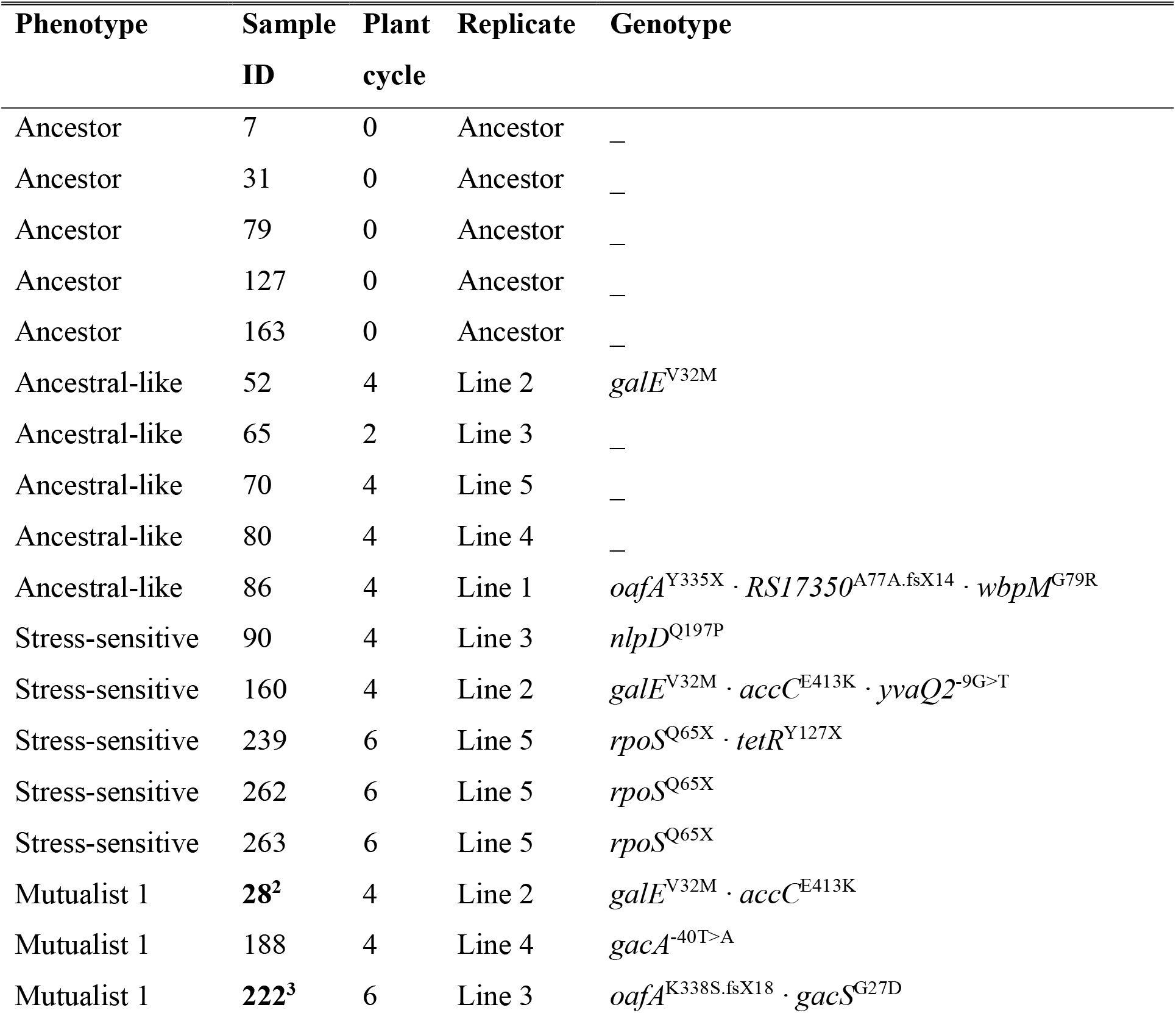
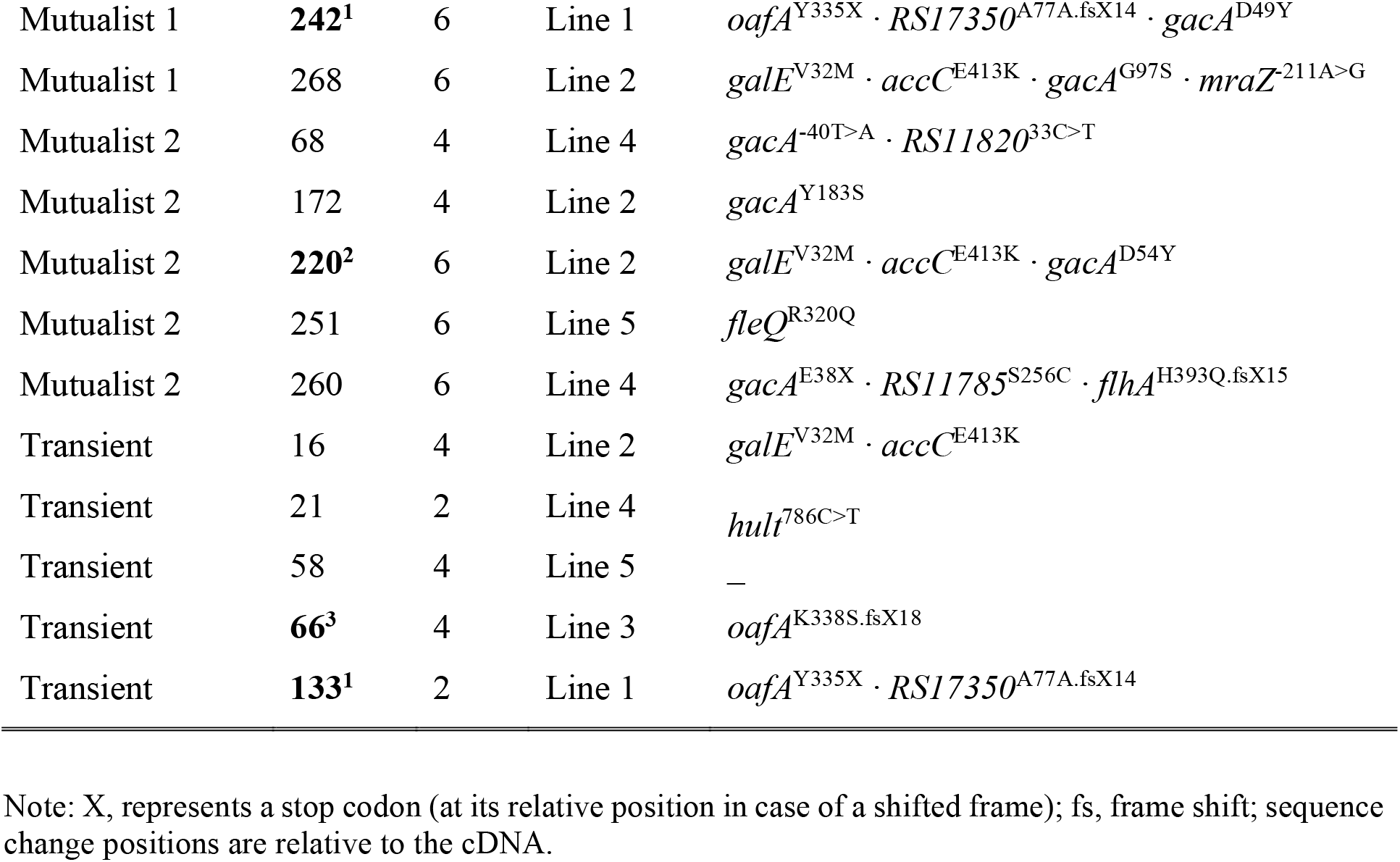
Description of 5 ancestral and 25 evolved *Pseudomonas protegens* CHA0 isolates, which were included to phenotyping and genotyping assays. “Sample ID” is the unique identifier of each isolate and five isolates were selected from each phenotype class. The “Plant cycle” column refers to the plant growth cycle from which the isolate was collected (see Figure S1) and the “Replicate” column refers to the independent plant replicate selection line. Mutated genes were identified using whole genome re-sequencing. On average, each evolved isolate encompassed 2-3 mutations relative to the ancestral sequence that are typically non-synonymous in nature, *i.e.* they directly affect predicted protein sequence and/or protein length. The same Sample ID superscript numbers shown on bold (133 and 242, 66 and 222, 28 and 220) indicate paired samples used in relative competition assays presented in Figure 5.

**Table S3.** Overview of the affected genes that are identified in the 25 evolved *Pseudomonas protegens* CHA0 isolates (See Table S2).

| *Gene* | *Product* | *Locus tag* | *DNA change* | *Effect* |
| --- | --- | --- | --- | --- |
| *accC* | Biotin carboxylase, acetyl-CoA carboxylase | PFLCHA0_RS03400 | c.1237 G>A | p.E413K |
| *flhA* | Flagellar biosynthesis protein FlhA | PFLCHA0_RS08490 | c.1154 T deleted | early stop |
| *gacA* | Response regulator GacA | PFLCHA0_RS17965 | c.548 A>C | p.Y183S |
| *gacA* | Response regulator GacA | PFLCHA0_RS17965 | c.289 G>A | p.G97S |
| *gacA* | Response regulator GacA | PFLCHA0_RS17965 | c.160 G>T | p.D54Y |
| *gacA* | Response regulator GacA | PFLCHA0_RS17965 | c.145 G>T | p.D49Y |
| *gacA* | Response regulator GacA | PFLCHA0_RS17965 | c.112 G>T | p.E38* |
| *gacA* | Response regulator GacA | PFLCHA0_RS17965 | -40 T>A | promoter |
| *galE* | UDP-glucose 4-epimerase | PFLCHA0_RS09920 | c.94 G>A | p.V32M |
| *RS11785* | LysR family transcriptional regulator | PFLCHA0_RS11785 | c.766 A>T | p.S256C |
| *hutI* | Imidazolonepropionase | PFLCHA0_RS02080 | c.786 C>T | synonymous |
| *RS11820* | PaaI family thioesterase | PFLCHA0_RS11820 | c.33 C>T | synonymous |
| *yvaQ2* | Methyl-accepting chemotaxis protein | PFLCHA0_RS13000 | -9bp G>T | promoter |
| *mraZ* | Transcriptional regulator mraZ | PFLCHA0_RS25175 | -211bp A>G | promoter |
| *nlpD* | Lipoprotein nlpD/lppB/LysM domain-containing protein | PFLCHA0_RS31060 | c.590 A>C | p.Q197P |
| *fleQ* | Sigma-54-dependent Fis family transcriptional regulator | PFLCHA0_RS08340 | c.959 G>A | p.R320Q |
| *oafA* | O-acetyltransferase OatA | PFLCHA0_RS09890 | c.1005 C>A | p.Y335* |
| *oafA* | O-acetyltransferase OatA | PFLCHA0_RS09890 | c.1009 A deleted | early stop |
| *wbpM* | Polysaccharide biosynthesis protein/NDP-sugar<br>epimerase | PFLCHA0_RS21855 | c.235 G>C | p.G79R |
| *rpoS* | RNA polymerase sigma factor RpoS | PFLCHA0_RS06125 | c.193 C>T | p.Q65* |
| *RS17350* | Methyltransferase domain-containing protein | PFLCHA0_RS17350 | c.116 C deleted | early stop |

**Table S4.**
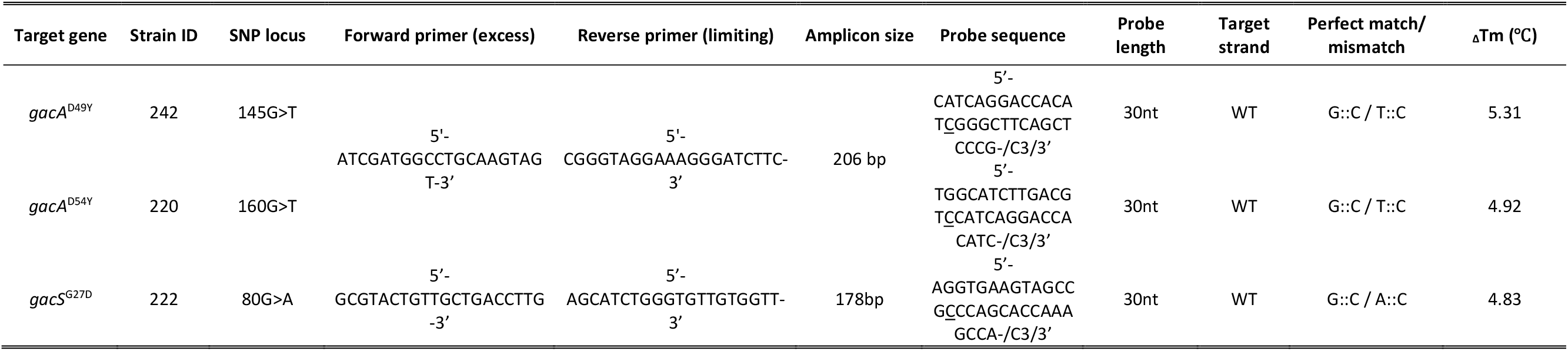
Primers and probes used for high-resolution melting (HRM) analysis. For the two *gacA* mutants the same set of primers was used. Underlined bases indicate the position of the single nucleotide point (SNP) mutations within the probe sequences. ΔTm (℃) indicates the melting temperature difference between WT-probe duplex and mutant-probe duplex.

**Supplementary dataset 1:** Sheet 1: Summary table for the set of fourteen phenotypic traits for the 256 characterised isolates used for K-mean clustering. Sheet 2: Carbon use data for the 256 characterised isolates. In both sheets, each line corresponds to one isolate and each column to one specific trait. See material and methods for a detailled description of experimental procedures.

**Supplementary dataset 2:** Sheet 1: Recapitulation of the origin (replicate line), time point and phenotype of each of the 30 isolates tested in details for their interactions with the host plant. Sheet 2: Summary table for interactions between each of the 30 isolates tested for plant growth, including effect on plant performance and induced GUS expression. Sheet 3: Scopoletin sensitivity. See material and methods for a detailled description of experimental procedures.

## Notes

### Competing Interest Statement

The authors have declared no competing interest.

